# Prickle2 regulates apical junction remodeling and tissue fluidity during vertebrate neurulation

**DOI:** 10.1101/2024.07.02.601771

**Authors:** Miho Matsuda, Sergei Y. Sokol

**Affiliations:** Department of Cell, Developmental and Regenerative Biology, Icahn School of Medicine at Mount Sinai, New York

**Author notes:** Correspondence: Miho Matsuda, Ph. D., Sergei Y. Sokol, Ph. D.

**Keywords:** Epithelial morphogenesis, neural tube closure, Xenopus embryo, planar cell polarity Adherens junctions, actomyosin

## Abstract

The process of folding the flat neuroectoderm into an elongated neural tube depends on tissue fluidity, a property that allows epithelial deformation while preserving tissue integrity. Neural tube folding also requires the planar cell polarity (PCP) pathway. Here, we report that Prickle2 (Pk2), a core PCP component, increases tissue fluidity by promoting the remodeling of apical junctions (AJs) in *Xenopus* embryos. This Pk2 activity is mediated by the unique evolutionarily conserved Ser-Thr-rich region (STR) in the carboxyterminal half of the protein. Mechanistically, the effects of Pk2 require Rac1 and are accompanied by increased cadherin dynamics and destabilization of tricellular junctions, the hotspots of AJ remodeling. Notably, Pk2 depletion leads to the accumulation of mediolaterally oriented cells in the neuroectoderm, whereas the overexpression of Pk2 or Pk1 containing the Pk2-derived STR promotes cell elongation along the anteroposterior axis. We propose that Pk2-dependent regulation of tissue fluidity contributes to anteroposterior tissue elongation in response to extrinsic cues.

## Introduction

Epithelial tissue fluidity is defined as the ability of epithelia to deform in response to stimuli while preserving structural integrity (Atia et al., 2021; Farhadifar et al., 2007; Guillot and Lecuit, 2013; Mongera et al., 2018; Park and Fredberg, 2016; Wang et al., 2020). In developing vertebrate embryos, an initially flat embryonic epithelium has a high potential to deform. In response to physical or chemical cues, individual epithelial cells change their shapes and neighbors in a highly organized anisotropic manner, leading to the formation of tissues or organs. Directional apical junction (AJ) remodeling is a key regulator of epithelial tissue deformation, however the underlying mechanisms are largely unknown (David et al., 2014; Founounou et al., 2021; Kim et al., 2021; Kuriyama et al., 2014).

The *Xenopus* ectoderm is a good model to study tissue fluidity during morphogenesis. At the onset of neurulation, the neuroectoderm is “fluid-like” with frequent cell intercalation and apical constriction, required for neural tube closure (NTC) (Baldwin et al., 2022b; Christodoulou and Skourides, 2022; Matsuda et al., 2023; Williams et al., 2014). In contrast, the non-neural ectoderm (future epidermis) is “solid-like”, characterized by hexagonal packing and a low rate of cell intercalation. Notably, epithelia can reacquire ‘fluidity’ during morphogenetic changes, allowing further deformation (Bocanegra-Moreno et al., 2023; Classen et al., 2005; Lawton et al., 2013; Mao and Wickstrom, 2024; Mitchel et al., 2020; Tetley and Mao, 2018). For instance, the reduced ‘fluidity’ of the embryonic ectoderm at the end of gastrulation is replaced by increased cell–cell rearrangements and apical domain heterogeneity during neurulation. Therefore, the spatiotemporal regulation of tissue and AJ fluidity is a hallmark of epithelial morphogenesis.

The planar cell polarity (PCP) pathway regulates the directionality of collective cell movements during morphogenesis. Originally defined in *Drosophila* genetic studies, core PCP components coordinate cell orientation in the tissue plane (Goodrich and Strutt, 2011; Humphries and Mlodzik, 2018; Peng and Axelrod, 2012). PCP is believed to be maintained by the feedback regulation of two complementary core protein complexes, Celsr-Vangl-Prickle (Pk) and Celsr-Frizzled (Fz)-Dishevelled (Dvl), which are distributed to the opposite side of cell junctions (Aw and Devenport, 2017; Davey and Moens, 2017; Peng and Axelrod, 2012). In the vertebrate mesoderm, the core PCP pathway promotes mediolateral cell intercalation during convergent extension (CE) movements, leading to embryonic axis elongation (Gray et al., 2011; Keller, 2002; Keller, 2012; Sokol, 1996; Wallingford et al., 2002). Besides CE defects, misregulated PCP signaling results in neural tube abnormalities (Colas and Schoenwolf, 2001; Henderson et al., 2018; Montcouquiol et al., 2003; Williams et al., 2014; Ybot-Gonzalez et al., 2007) that may involve defects in apical constriction (Ossipova et al., 2015b) and radial cell intercalations (Ossipova et al., 2015a). Altogether, these studies suggest that PCP signaling is relevant to tissue fluidity control.

The core PCP components are functionally linked to the actomyosin cytoskeleton. The Fz-Dvl complex binds a RhoGEF and Daam1, a formin family protein that activates RhoA in the cell cortex (Habas et al., 2001; Nishimura et al., 2012). The Fz-Dvl complex may, therefore, promote the formation of supracellular actomyosin cables that bend the neural plate (McGreevy et al., 2015; Nishimura et al., 2012; Roper, 2013). The Vangl-Pk complex is enriched at the anterior cell edge in various vertebrate models (Butler and Wallingford, 2018; Ciruna et al., 2006; Devenport and Fuchs, 2008; Mancini et al., 2021; Ossipova et al., 2015c; Yin et al., 2008). However, the cellular targets of Vangl and Pk mediating tissue deformation largely remain to be identified.

In this study, we investigate the role of Prickle2 (Pk2) in AJ remodeling and tissue fluidity control during neurulation. Consistent with its presumed function as a core PCP protein, Pk2 is polarized at the anterior AJ of *Xenopus* neuroepithelial cells, and Pk2 morphants exhibit NTC defects (Butler and Wallingford, 2018). The same study showed a correlation between Pk2 and actomyosin enrichment in shrinking AJs; however, the underlying mechanism and causation remain unclear. Here we show that Pk2 knockdown attenuates AJ elongation and shrinkage in the neuroectoderm. Conversely, Pk2 overexpression in the gastrula ectoderm inhibits hexagonal cell packing, characteristic of the cells at the end of gastrulation. This activity of Pk2 is unique and involves an evolutionarily conserved Ser/Thr-rich region of the protein. It also requires Rac1 activation and is accompanied by increased dynamics of cadherin and actomyosin at AJs. Importantly, the effects of Pk2 on AJ remodeling are anisotropic in the neuroectoderm and induce preferential elongation of cells along the anteroposterior axis. We propose that Pk2 increases epithelial tissue fluidity, leading to an enhanced response of the epithelia to anisotropic cues from the environment.

## Results

### Pk2 depletion decreases the rate of AJ remodeling in the neuroectoderm

We assessed the role of Pk2 in AJ remodeling and tissue fluidity in *Xenopus* neuroectoderm (NE), a “fluid-like” epithelium characterized by intensive AJ remodeling, cell intercalation and apical domain variability (Baldwin et al., 2022a; Christodoulou and Skourides, 2022; Matsuda et al., 2023; Williams et al., 2014). A previously characterized splicing-blocking morpholino (MO) (Butler and Wallingford, 2015; Butler and Wallingford, 2018; Devitt et al., 2024; Shindo et al., 2019) was co-injected with a lineage tracer into one dorsal blastomere at the 4– to 8-cell stage, to unilaterally deplete Pk2 in the NE (Fig. 1A). The unilateral knockdown minimizes anteroposterior (AP) axis elongation defects and allows to compare behaviors of the morphant and wild type NE cells in the same embryo (Fig. 1A, left to right). We assessed changes in AJ length, orientation, and cell‒cell rearrangements in the posterior NE close to the brain‒spinal cord border, where approximately 25% of cells intercalate during neurulation (Butler and Wallingford, 2018), leading to the replacement of the mediolaterally oriented (ML) junctions with anteroposteriorly oriented (AP) junctions.

**Figure 1.**
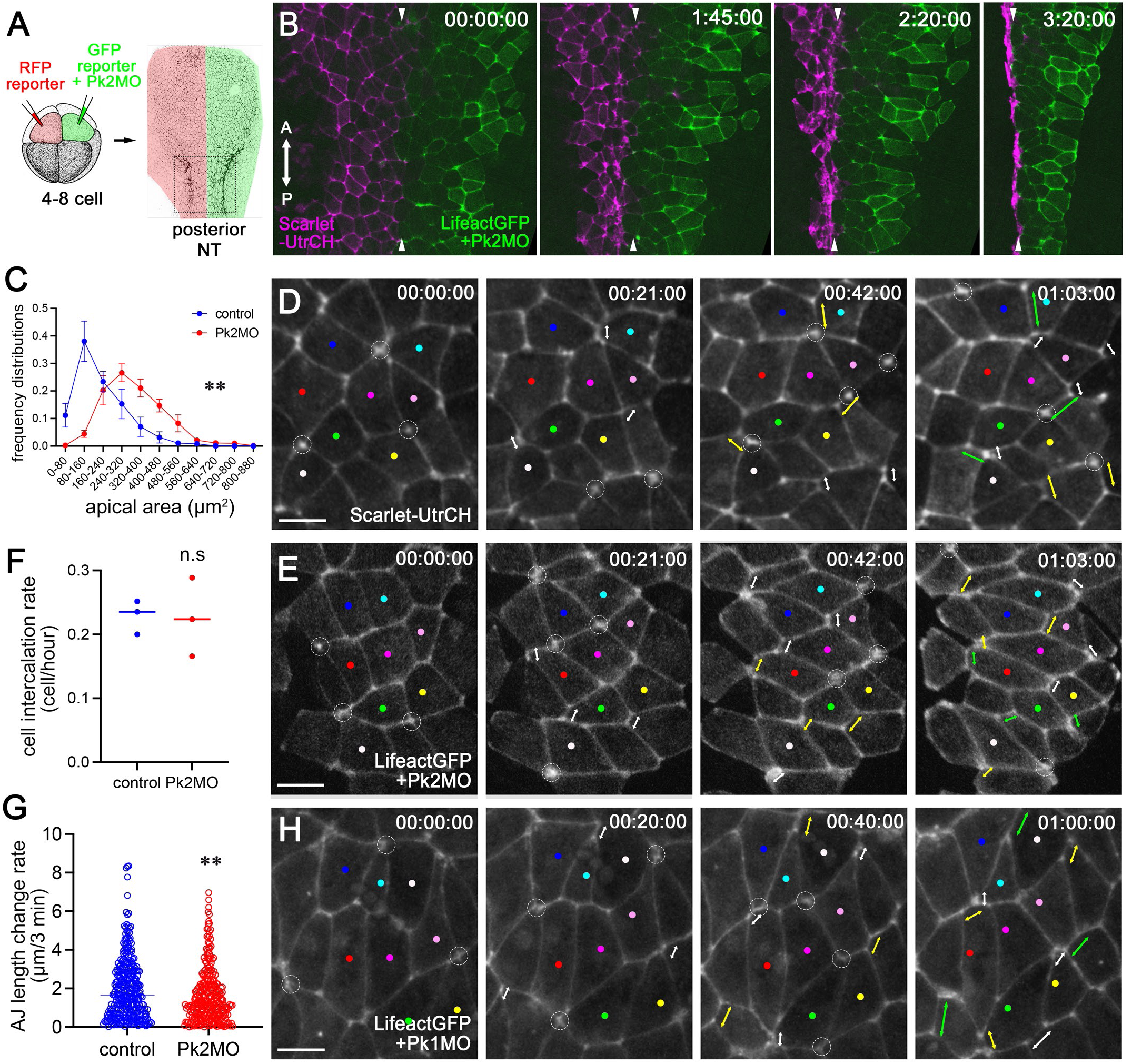
**Pk2 knockdown in the *Xenopus* neuroectoderm attenuates AJ remodeling** (A) Experimental scheme. Pk2 MO was injected with myrGFP or LifeactGFP RNA into one dorsal animal blastomere (green) at the 4-8-cell stage. myrRFP or Scarlet-UtrCH RNA was injected into the other dorsal animal blastomere (red) as a control. The superficial neuroectoderm (NE) of the posterior NT in stage 14-18 embryos was imaged near the brain‒ spinal cord border. (B) Representative images of the control (Scarlet-UtrCH) and Pk2 MO (LifeactGFP) NEs from Video 1. Imaging started at stage 15, with 5 min intervals and a duration of 3 hrs 40 min. Arrowheads indicate the midline. (C) Frequency distribution histogram of apical domain size. 3 embryos per group. Stage 16. (D, E) Representative control (D) and Pk2 KD (E) images of stage 14-16 NEs from Video 2. The colored dots label individual cells. White circles indicate the initiation of the T1 transition. The white, yellow and green double-headed arrows indicate the progressive elongation in newly formed AJs after T1 transitions. (F) Frequency of cells that entered the T1 transition in the control and Pk2-KD NEs. Dots represent the average frequency of T1 transition in the individual NEs. Three each. (G) Rate of AJ length change in the control and Pk2 KD NEs. Both AJ elongation and shortening are included. F, G: n=312 cells (control); n=285 cells (Pk2 KD). The control and Pk2 KD side of the NE in the same time-lapse imaging field were compared in individual embryos. (H) Representative still images of Pk1-KD NEs from Video 5. Stage 14-16. 5 min intervals. Duration, 1 hr 45 min. A: anterior. P: posterior. Student’s t test. **p<0.01. Scale bars: 20 μm.

Pk2 knockdown (KD) disrupted NTC (Fig. 1B, Video 1) as reported previously (Butler and Wallingford, 2018). Time-lapse imaging revealed significant defects in apical domain size reduction in Pk2 KD cells (Fig. 1C). Pk2 KD cells initiated T1 transitions, the exchange of neighbors among four cells, with the same frequency as the control cells during a <2-hour time lapse imaging (white circles in Fig. 1D and 1E, Video 2). The orientation of the T1 transitions was also normal (a total of 277/277 ML to AP intercalation events in three Pk2 KD embryos), suggesting that Pk2 is not essential for T1 transition. However, the elongation of newly formed AP junctions after T1 transitions was severely attenuated (double arrowheads in white, yellow and green track the progressive elongation of newly formed AP junctions in Fig. 1D and E). AJ elongation defects in Pk2 MO cells were not limited to AP junctions that were newly formed after T1 transition. Tracking of randomly selected junctions in the NE showed that Pk2 KD reduced the rate of both elongation and shrinkage of AJs (Fig. 1G). These results suggest that Pk2 is required for efficient AJ elongation and shrinkage in the NE.

### Posterior NE cells do not extend along the AP axis after Pk2 depletion

Another noticeable change in the Pk2 MO cells was their orientation in the NE. Wild-type cells were predominantly ML-oriented in the early neurula and progressively acquired AP orientation during neural fold formation (myrRFP in Fig. 2A and Video 3, Fig. 2B). Pk2-depleted cells retained the ML orientation in the folding neural plate (myrGFP+Pk2MO in Fig. 2A and Video 3, Fig. 2C). We confirmed this observation by phalloidin staining of fixed embryos (Fig. S1). Although Pk1 depletion also disrupted NTC (Video 4), Pk1-depleted NE cells oriented along the AP axis just like the wild-type cells (Fig. 1H, Video 5). These results indicate that the observed cell orientation defect is specific to Pk2.

**Figure 2.**
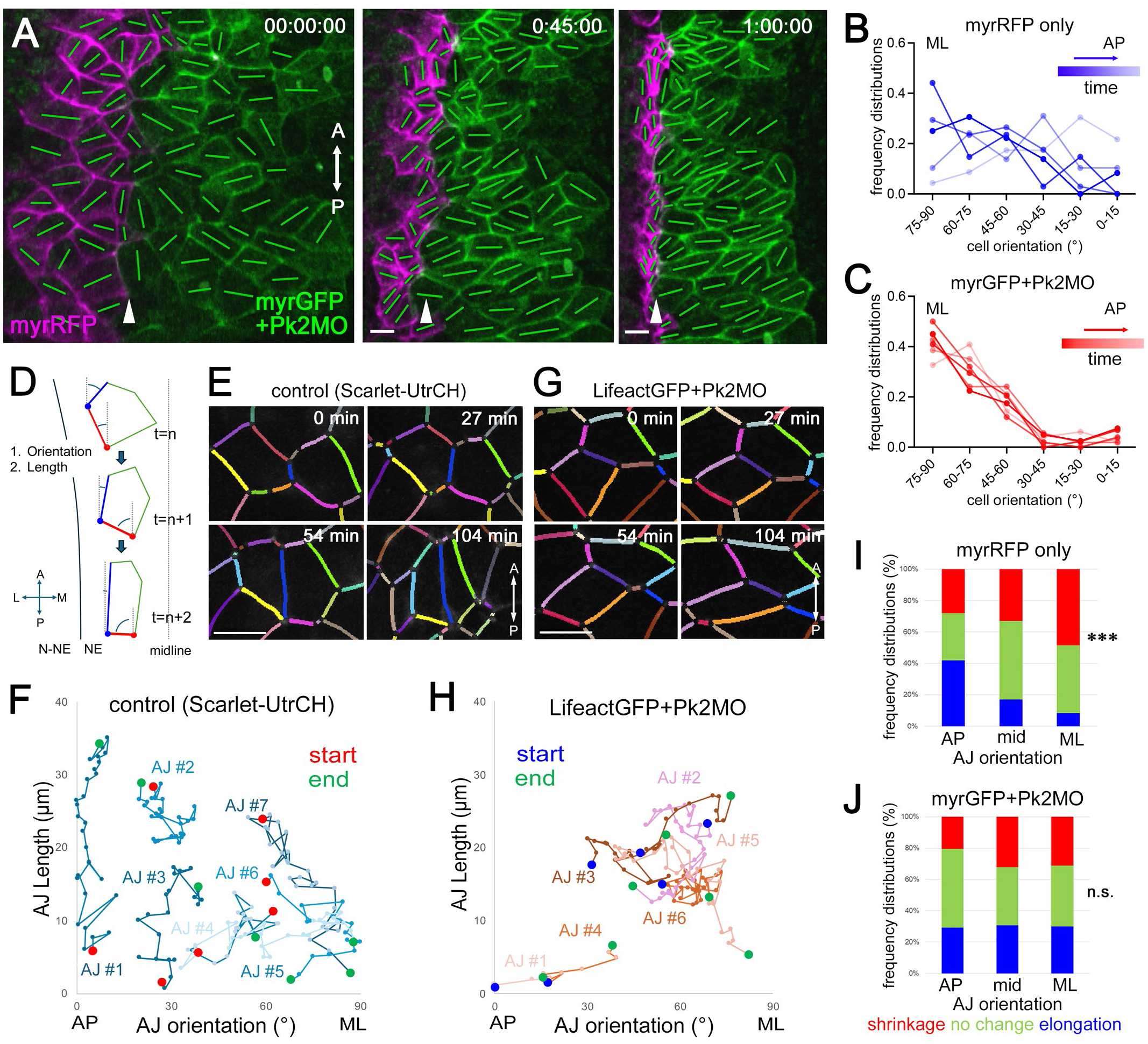
**Effects of Pk2 depletion on AJ orientation in the neuroectoderm** (A) Still images from time-lapse imaging of the NEs, including control (myrRFP) and Pk2 KD (myrGFP) cells (Video 4, 1 hour 12 min, 3 min intervals). Stage 14-16. The green lines indicate the cell orientation in individual cells. A: anterior. P: posterior. The white arrowhead indicates the presumed midline. (B, C) A representative frequency distribution histogram of cell orientation in the control (B) and Pk2 KD (C) NEs. Circular angles 0 and 90 correspond to the anteroposterior (AP) and mediolateral (ML) axes, respectively. The darker colors represent the NE at earlier time points of the time-lapse imaging. (D) A scheme of the AJ quantification method used in Fig. 2E-J and Fig. 10. After time-lapse imaging, individual AJs were tracked, and their orientation and length were quantified. (E) Representative images of AJ tracking in the wild-type NEs. (F) A representative 2D plot of AJ length and orientation changes in the wild-type NE. X-axis: AJ orientation. Y-axis: AJ length. Individual blue dots represent the orientation and length of single AJs, which were tracked for 1 hour and 28 min (connected by the blue lines). Red and green dots indicate the orientation and length of AJs at the start and end of tracking, respectively. The plot includes the tracking and quantification of 7 AJs. (G, H) Representative images of AJ tracking and quantification in Pk2-KD NEs. Blue and green dots indicate the orientation and length of AJs at the start and end of tracking, respectively. The 2-D plot includes 6 AJs. (I, J) Quantification of changes in AJ length in the control NE (I) and the Pk2 MO NE (J) groups. Changes in AJ length are defined as follows: elongation (blue, ≥0.8 μm/3 min), shrinkage (red, ≤-0.8 μm/3 min), or no change (green, –0.8< and <0.8 μm/3 min). AJs are grouped, based on orientation, into AP (≤30° from the AP axis), ML (≤30° from the ML axis), and mid (30-45° from either the AP or ML axes). The graphs include data obtained from tracking 17 AJs (control) and 15 AJs (Pk2 KD) for 1 h and 28 min. The control and Pk2 KD side of the NE in the same time-lapse imaging field were compared. The total number of data points: n=353 (control) and n=595 (Pk2 KD). Chi-square test for I and J. ***p<0.001. n.s., not significant. Scale bars: 20 μm.

We tracked changes in AJ length and orientation to further define defects in the Pk2 KD NE (Fig. 2D). In the wild-type NE, average AP junctions were longer than ML junctions (Fig. 2E, F). Individual AP junctions preferentially elongated as NTC progressed, while ML junctions shrank (Fig. 2E, F, I), consistent with previous reports (Baldwin et al., 2022a; Christodoulou and Skourides, 2022). In addition, individual AJs changed their orientation (Fig. 2D-F), reorienting cells from ML to AP (Fig. 2B). By contrast, in the Pk2-depleted NE, average ML junctions were longer than AP junctions (Fig. 2G, H). There was no significant difference in the proportion of AJ length change between AP and ML junctions (Fig. 2J). We confirmed these findings by time-lapse imaging and AJ tracking in another embryo (Fig. S1D, E). These results suggest that Pk2 is required for the orientation-dependent AJ behavior control.

### Pk2 overexpression inhibits hexagonal cell packing in *Xenopus* ectoderm

We next asked whether this Pk2 activity in AJ remodeling is specific for the NE. Flag-Pk2 RNA was injected into the superficial gastrula ectoderm (Fig. 3A) that has well-organized AJs and apical‒basal polarity (Cardellini et al., 2007; Fesenko et al., 2000). Importantly, tissue fluidity changes in the ectoderm during gastrulation (myrGFP only in Video 6): At the early gastrula stage (stage 10, 00:00 h in Fig. 3B), ectodermal cells were diverse in size and shape (Fig. 3G, J) with curved cell‒cell boundaries (Fig. 3H, I). In the late gastrula stage (stage 11.5, 04:55 h in Fig. 3B), the cells became more uniform in size (Fig. 3G), hexagonal in shape (Fig. 3J) and with linear cell-cell borders (Fig. 3I), consistent with increased hexagonal packing (Classen et al., 2005; Farhadifar et al., 2007).

**Figure 3.**
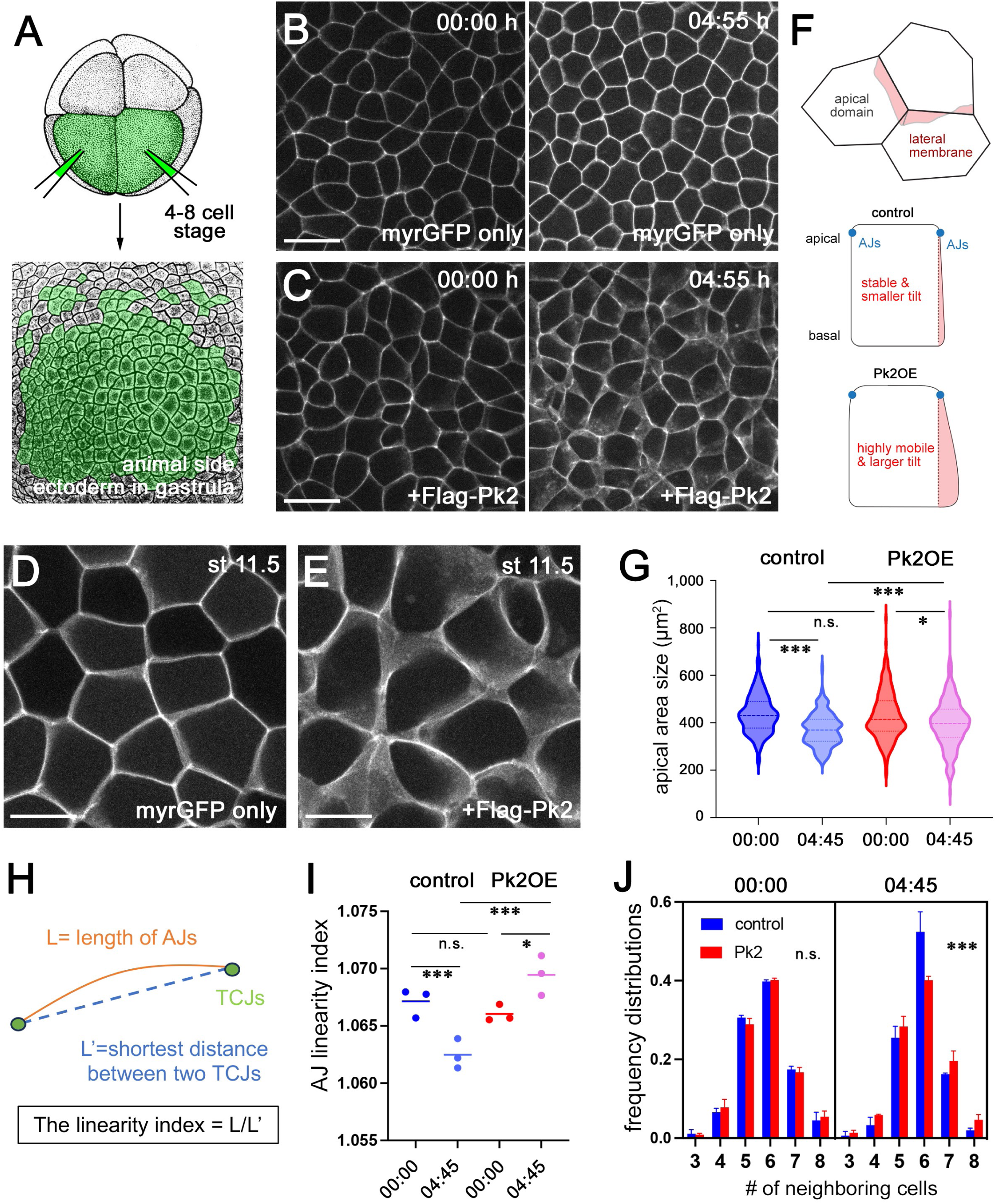
**Pk2 inhibits hexagonal packing in the gastrula ectoderm** (A) Scheme of the experiment. Flag-Pk2 mRNA was coinjected with myrGFP mRNA into two ventral blastomeres at the 4-8-cell stage. Time-lapse imaging of the animal superficial ectoderm was initiated at stage 10. (B, C) Gastrula ectoderm expressing myrGFP only (B) and myrGFP with Flag-Pk2 (C). Snap-shot images from Video 6. (D-F) Higher-resolution images of ectoderms expressing myrGFP only (D) or with Flag-Pk2 (E) at stage 11.5. Snap images from Video 7. Albino embryos were used to assess the dynamics of myrGFP fluorescence signals at the lateral membrane, as shown in the schematic diagram (F). (G-J) Quantification of hexagonal cell packing. Images obtained from the pigmented wild-type embryos were used for segmentation. Cell packing was assessed by apical domain size heterogeneity (G), AJ linearity (I) and the number of neighboring cells (J). Three embryos were used per group. (H) Schematic of the quantification of AJ linearity. AJ linearity was determined by the actual length of the AJs divided by the shortest distance between two vertices (L/L’). (I) Individual dots represent the average AJ linearity in each embryo. The lines correspond to the means of three embryos. (J) The frequency distribution histogram of the number of neighboring cells. The total numbers of cells scored in G-J: 576 cells and 1,127 AJs (myrGFP control, 00:00 h); 672 cells and 1,169 AJs (myrGFP control, 4:55 h); 724 cells and 1,360 AJs (myrGFP+F-Pk2, 00:00 h); and 764 cells and 1,355 AJs (myrGFP+F-Pk2, 4:55 h). The Kruskal‒Wallis test for G. Student’s t-test for I. Chi-square test for J. *<0.05, **p<0.01, ***p<0.001. n.s., not significant. Scale bars: 50 μm (B, C), 25 μm (D, E).

In the Pk2-overexpressing (OE) ectoderm in early gastrula (stage 10, 00:00 h in Fig. 3C), the apical domain size variability (Fig. 3G), AJ linearity (Fig. 3I) and number of neighbors (Fig. 3J) were comparable to the corresponding parameters in wild-type ectoderm. However, as gastrulation progressed, Pk2 OE cells continued to exhibit variable apical domains (04:55 h in Fig. 3C, G, I, J, and myrGFP+Flag-Pk2 in Video 7). Pk2 OE cells also had increased lateral membrane dynamics (Fig. 3D-F, and Video 8) and more frequent cell intercalations (Fig. 4A-C), as compared with wild-type control cells. These findings suggest that Pk2 inhibits hexagonal packing in the gastrula ectoderm. This is complementary to Pk2 morphant phenotype in the NE.

**Figure 4.**
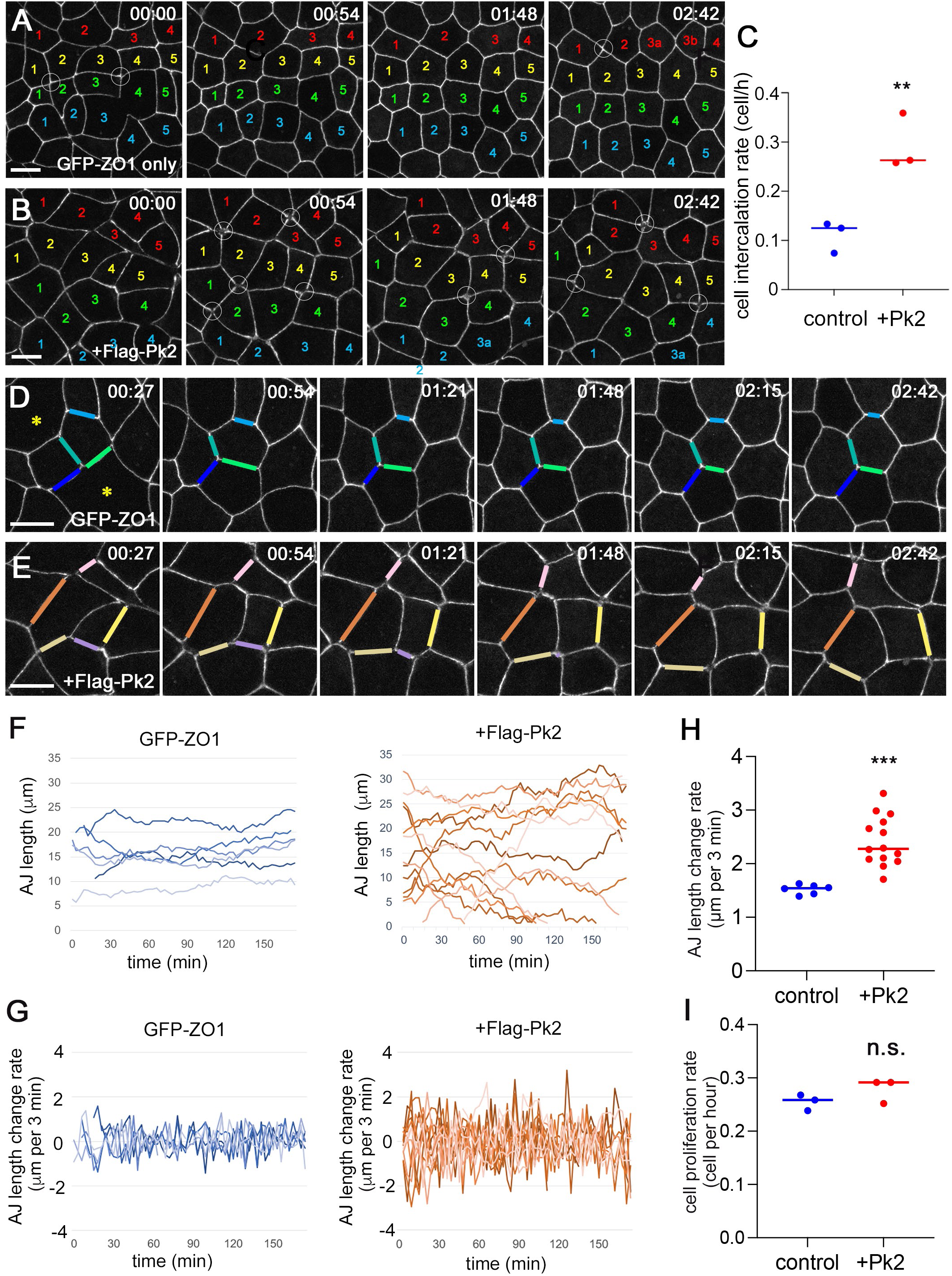
**Pk2 promotes AJ remodeling and cell intercalation in the gastrula ectoderm** (A, B) Representative snapshot images of *Xenopus* gastrula ectoderm. GFP-ZO1 only (A) and GFP-ZO1 with Flag-Pk2 (B). Time-lapse imaging was initiated at stage 11. The white circles indicate the initiation of T1-cell intercalation. Asterisks show cells which divided during the time-lapse imaging. (C) The rate of cell intercalation in the control or Pk2 ectoderm. Each data point is the cumulative sum from one embryo. Three embryos total. (D, E) Representative images of AJs in control GFP-ZO1 only (D) or with Flag-Pk2 OE (E) ectoderms. (F) Changes in AJ length. The plots included 6 AJs (control) and 14 AJs (+Flag-Pk2). (G) Rate of change in AJ length, shown in F. (H) The average rate of change in AJ length. Individual dots represent the average AJ length changes in individual AJs, as shown in G. (I) Cell proliferation rate in stage 10-11.5 embryos expressing GFP-ZO1 only or GFP-ZO1 with Flag-Pk2. Three embryos per group. The total numbers of cells in C and I: n=73 (control); n=124 (Flag-Pk2). The total number of AJ length changes in F-H: n=374 (control); n=412 (Flag-Pk2). The median line is indicated in H and I. Student’s unpaired t test was used for C, H and I. **p<0.01, ***p<0.001, n.s., not significant. Scale bars: 20 μm.

### Pk2 promotes AJ remodeling in the ectoderm

Dynamic remodeling of AJs increases tissue fluidity (Pinheiro and Bellaiche, 2018). To assess AJ dynamics in the Pk2-expressing ectoderm, we tracked the rate of AJ elongation and shrinkage in nonmitotic cells (Fig. 4D-H) and the rate of AJ elongation in daughter cells after cytokinesis (Fig. S2). In both cases, Pk2 OE accelerated AJ elongation and shrinkage (Figs. 4F-H, S2C). The mitotic rate was similar between Pk2 OE and control ectoderm (Fig. 4I), excluding the potential effect of cell division on tissue fluidity (Devany et al., 2021; Firmino et al., 2016; Matoz-Fernandez et al., 2017; Ranft et al., 2010). These results indicate that Pk2 increases tissue fluidity by promoting AJ remodeling in gastrula ectoderm.

Increased cadherin turnover has been reported in AJs under dynamic remodeling (Kowalczyk and Nanes, 2012). Time-lapse imaging of YFP-C-cadherin (Video 8) showed that the distribution of C-cadherin in the Pk2 OE ectoderm was less uniform at bicellular AJs (BJs) (Fig. 5A-C). C-cadherin puncta in the cytoplasm increased (arrowheads in Fig. 5A, B), suggesting that C-cadherin at AJs is destabilized in the Pk2 OE ectoderm. Tricellular junctions (TCJs) are hotspots of AJ remodeling and actomyosin tension, where AJ and actomyosin components accumulate (Bosveld et al., 2018; Finegan et al., 2019; Letizia et al., 2019; Uechi and Kuranaga, 2019; Vanderleest et al., 2018). Pk2 OE reduced C-cadherin abundance at TCJs (Fig. 5E) and increased C-cadherin fluctuations at TCJs over time (Fig. 5D, F, G). Increased fluctuation of E-cadherin has been reported in mobile AJs (Huebner et al., 2021; Vanderleest et al., 2018). These results suggest that the TCJs in the Pk2 OE ectoderm are more mobile than those in the wild-type ectoderm.

**Figure 5.**
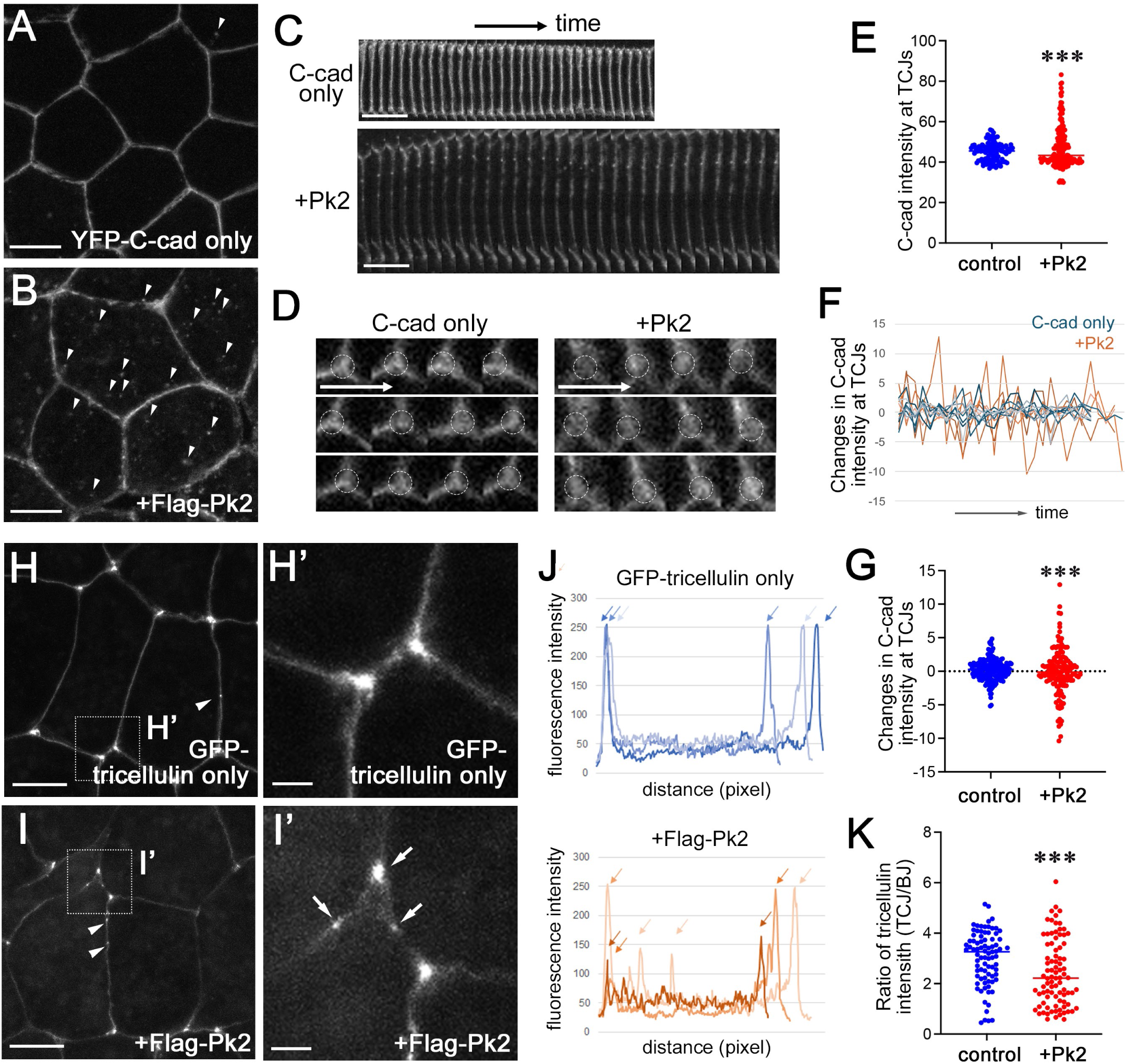
**Destabilization of AJs and TCJs in Pk2 OE ectoderm** (A, B) Representative images of YFP-C-cadherin in the control (A) or Pk2-expressing (B) ectoderm. Arrowheads indicate C-cadherin puncta in the cytoplasm. Stage 11.5. (C, D) Representative kymographs of YFP-C-cadherin at bicellular junctions (BJs) (C) and tricellular junctions (TCJs) (D) from Video 8 with durations of 14 min 30 sec, and 30 second time intervals. Single *z*-plane images. (E) Quantification of fluorescence intensity of YFP-C-cadherin at TCJs.186 TCJs (control) and 151 TCJs (+Flag-Pk2) from three embryos. (F, G) Representative changes in YFP-C-cadherin fluorescence intensity at TCJs from Video 8. 7 TCJs (control) and 6 TCJs (+Flag-Pk2) are shown. (H-I’) Representative images of GFP-tricellulin in stage 11 control (H) and Flag-Pk2 (I) ectoderms. The square areas are enlarged in H’ and I’. The arrowheads in H and I indicate ectopic enrichment of GFP-tricellulin at the BJs. The arrows in I’ indicate disintegrated TCJs. (J) Fluorescence intensity plots of GFP-tricellulin from three independent AJs. Two vertices of individual AJs are indicated by red, blue and green arrowheads. (K) Quantification of GFP-tricellulin intensity at TCJs. The fluorescence intensity of GFP-tricellulin at individual TCJs was divided by the average fluorescence intensity of GFP-tricellulin at BJs. 83 TCJs (control) and 87 TCJs (Pk2). Three embryos per group. The Kolmogorov-Smirnov test was used for E and K. Chi-square test was used to quantify the categorized data in G. ***p<0.001. n.s. Scale bars: 20 μm (A-C, H, I), 10 μm (C), 5 μm (D), and 2 μm (H’, I’).

Furthermore, Pk2 OE reduced tricellulin at TCJs (Fig. 5H, I, K). Tricellulin is a TCJ-specific protein essential for junctional structural integrity and barrier function (Bosveld and Bellaiche, 2020; Ikenouchi et al., 2005). This was accompanied by increased tricellulin localization at BJs (arrowheads in Fig. 5H, I, arrows in Fig. 5J) and disruption of TCJs (arrows in Fig. 5I’). Non-muscle myosin II (NMII) and αE-catenin also reduced the accumulation at TCJs (Fig. S3), indicating a general disruption of TCJs. Taken together, these results suggest that TCJs are destabilized in the Pk2 OE ectoderm, a finding consistent with enhanced AJ remodeling.

### The Ser-Thr-rich (STR) region of Pk2 is required for Pk2-induced tissue fluidity increase

To determine the domains of Pk2 required for tissue fluidity increase, Pk2 deletion constructs (Fig. 6A) were overexpressed in the gastrula ectoderm. Tissue fluidity was assessed by AJ linearity (Fig. 6H), and the variability of junction length (Fig. 6I) and the number of neighboring cells (Fig. 6J). Three domains were required for the Pk2OE-induced increase in tissue fluidity. First, the C-terminal CAAX motif was required (Pk2ϕλC30 in Figs. 6G-J), a domain mediating the membrane localization of Prickle proteins (Fig. S4E)(Cho et al., 2015). Second, it required the Vangl-binding domain (VBD) (see Pk2ϕλVBD in Figs. 6F, H-J)(Butler and Wallingford, 2015; Jenny et al., 2003), even though Pk2ϕλVBD was localized at AJs (Fig. S4D) and Vangl2 did not increase tissue fluidity on its own (Figs. S5H, J-L). These observations suggest that Pk2 cooperates with Vangl2 at AJs for full activity.

**Figure 6.**
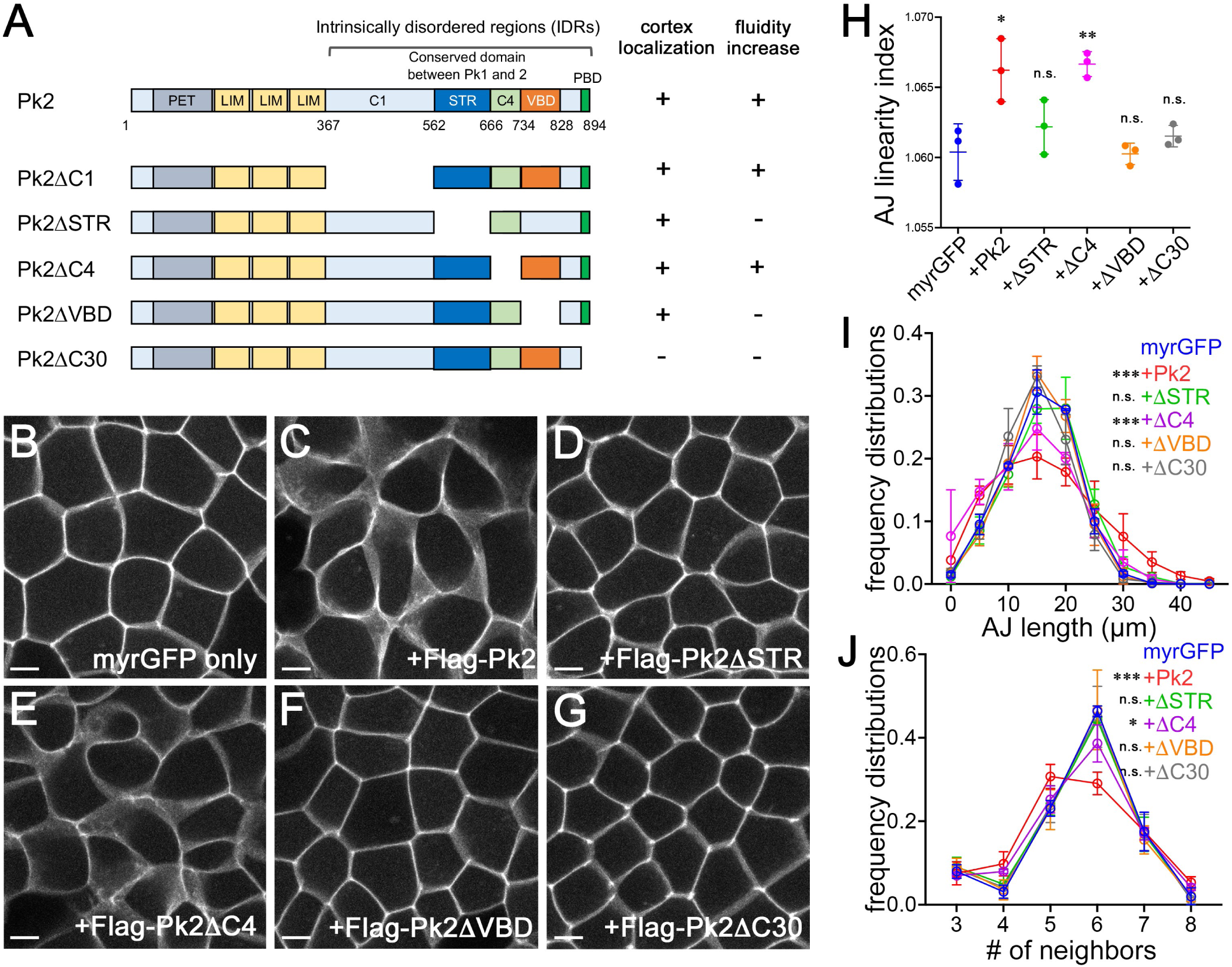
**The STR region is required for Pk2-mediated tissue fluidity control** (A) Scheme of Pk2 deletion mutant constructs. Pk2 contains one PET and three LIM domains in the N-terminal half and intrinsically disordered regions (IDRs) and a PDZ domain binding motif (PBD) in the C-terminal half. The IDRs include the Ser-Thr-rich region (STR) and the Vangl-binding domain (VBD). Pk2τιC30 lacks the C-terminal 30 amino acids, including the PBD. (B-G) Representative images of stage 11 ectoderms expressing myrGFP only (B) or myrGFP with Pk2 deletion mutants (C-G). Albino embryos were used to visualize the lateral domain. (H) AJ linearity was quantified as described in Fig. 3G. Individual dots represent the average AJ linearity of three embryos. (I) Frequency distribution histogram of AJ length. (J) Frequency distribution of the number of neighboring cells. Three embryos were used per condition. 345 cells and 1,052 AJs (myrGFP). 566 cells and 1,723 AJs (Pk2). 306 cells and 1,019 AJs (Pk2τιSTR). 429 cells and 1,305 AJs (Pk2τιC4). 384 cells and 1,150 AJs (Pk2τιVBD). 428 cells and 1,272 AJs (Pk2ι1C30). The statistical significance of differences was determined between control cells and cells expressing Pk2 deletion mutants. Student’s t test for H. The Kruskal‒ Wallis test for I. Chi-square test was used to quantify the categorized data for J. *<0.05, **p<0.01, ***p<0.001. n.s., not significant. Scale bars: 10 μm.

The third is the serine-threonine rich region (STR) (Pk2ϕλSTR in Fig. 6D, H-J), an evolutionarily conserved domain among vertebrate Pk1 and Pk2 (Fig. S5A, B) but absent in invertebrate Prickle and vertebrate Pk3. We confirmed that Pk2ϕλSTR localized at AJs in the gastrula ectoderm (Fig. S4B). Other regions of Pk2, such as C4, were not required for the AJ localization (Fig. S4C) or the increase in tissue fluidity (Fig. 6E, H-J).

### The STR region is responsible for functional differences between Pk1 and Pk2 in tissue fluidity control

Previous studies have implicated Pk1 in cell motility and F-actin reorganization (Carreira-Barbosa et al., 2003; Huang and Winklbauer, 2022; Tao et al., 2009; Veeman et al., 2003) through interactions with various GEFs and GAPs (Daulat et al., 2019; Zhang et al., 2016; Zhang and Wrana, 2018). ARHGAP21/23 binds Pk1 via a region including the STR and inhibits RhoA activation, resulting in disorganization of the actomyosin network and altered focal adhesion dynamics in cultured mammalian cells (Zhang et al., 2016). In our assays, Pk1 KD inhibited NTC (Fig. 1H), supporting its role in F-actin regulation and cell motility. However, Pk1 OE did not inhibit hexagonal epithelial packing (Figs. 7B, H, I, K, S5D) or reduce the localization of AJ or actomyosin components at TCJs (data not shown) in the late gastrula ectoderm. These results suggest that Pk1 and Pk2 modulate AJs or actomyosin via distinct mechanisms.

**Figure 7.**
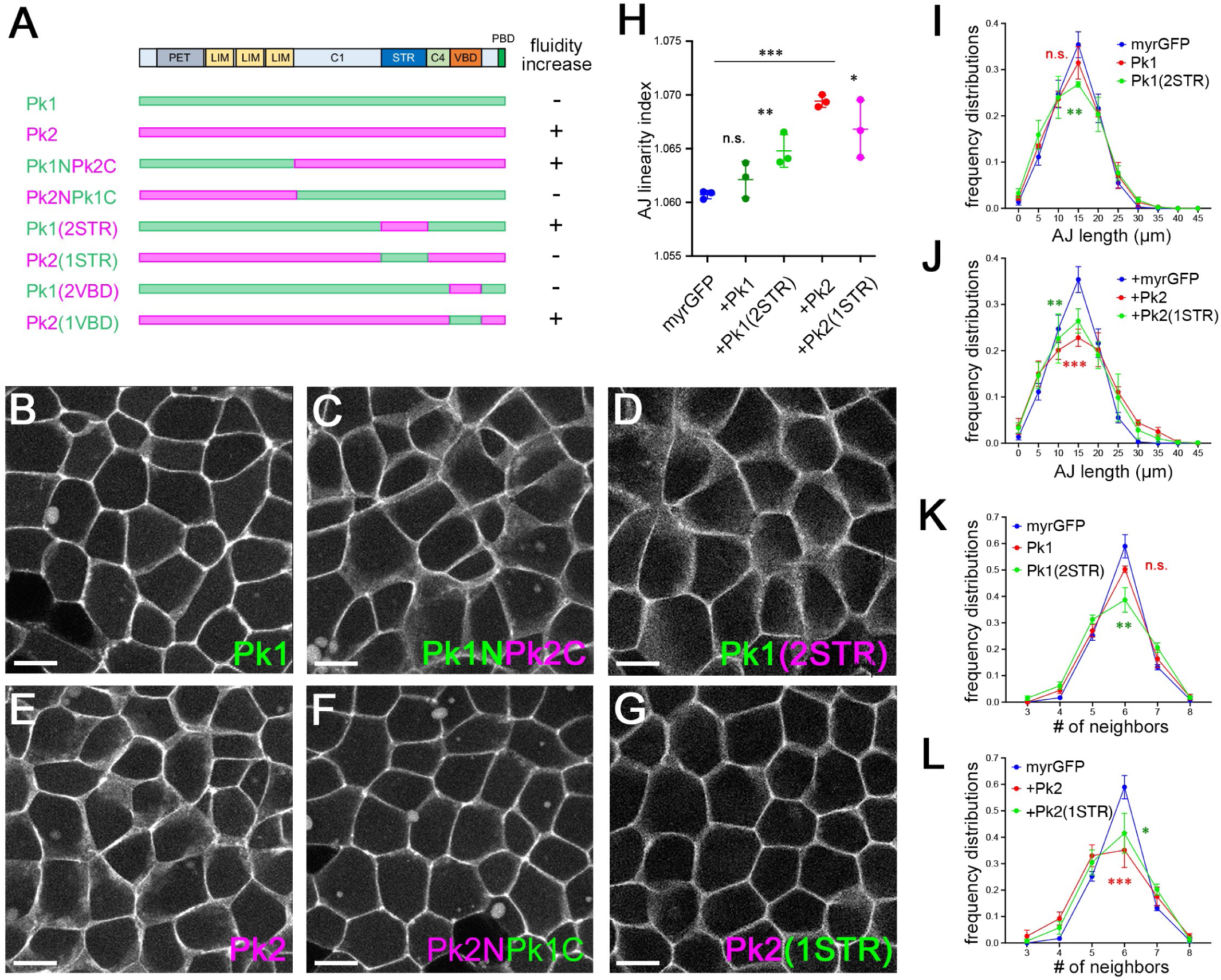
**The STR region is responsible for tissue fluidity control** (A) Schematic diagrams of the Pk1-Pk2 chimeric mutant constructs. (B-G) Representative images of stage 11 ectoderms expressing myrGFP with HA-Pk1 (B), Flag-Pk1NPk2C (C), HA-Pk1(2STR) (D), Flag-Pk2 (E), HA-Pk2NPk1C (F), and Flag-Pk2(1STR) (G). Albino embryos. (H-J) Tissue fluidity was evaluated by scoring AJ linearity (H), AJ length distribution (I), and the number of neighboring cells (J). Three embryos per group. 420 cells and 1,297 AJs (Pk1). 458 cells and 1,339 AJs (Pk1(2STR)). 372 cells and 1,110 AJs (Pk2). 399 cells and 1,228 AJs (Pk2(1STR)). Student’s t test (H), the Kruskal‒Wallis test (I, J) and chi‒square test (K, L) were used. *<0.05, **p<0.01, ***p<0.001. n.s., not significant. Scale bars: 20 μm.

To determine the domain responsible for the functional difference between Pk1 and Pk2, chimeric Pk1 and Pk2 proteins were constructed (Fig. 7A). Pk1N-Pk2C, a chimeric protein containing the N-terminal region of Pk1 and the C-terminal region of Pk2, increased tissue fluidity in the gastrula ectoderm (Fig. 7C), but Pk2N-Pk1C did not (Fig. 7F). These results suggest that the C-terminal region of Pk2 is sufficient to confer the tissue-fluidizing activity of Pk2 to Pk1. We next expressed Pk1 and Pk2 with swapped STRs (Fig. 7A) in the gastrula ectoderm. The insertion of the Pk1 STR domain into the Pk2 backbone (Pk2(1STR)) suppressed the tissue-fluidizing activity of Pk2 (Fig. 7G, H, J, L), whereas the complementary construct, Pk1(2STR), had the opposite effect (Fig. 7D, H, I, K). Notably, Pk3, which does not contain STR (Fig. S5A), did not affect ectoderm fluidity (Fig. S5G, I-K). The VBD swap did not alter tissue fluidity (data not shown). These results suggest that the STR is responsible for the functional difference between Pk1 and Pk2.

### Pk2 requires Rac1 to increase tissue fluidity in the gastrula ectoderm

Rho family small GTPases regulate AJs and actomyosin (Arnold et al., 2017; Citi et al., 2014; Heasman and Ridley, 2008). ARHGAP21 and ARHGAP23, GAPs for RhoA, bind the STR domain of Pk1 and inhibit RhoA activity (Zhang et al., 2016). Our attempt to test Pk2 binding to ARHGAP21/23 was not successful (data not shown). Alternatively, we coexpressed the dominant interfering forms of RhoA or Rac1 (DN-RhoA or DN-Rac1) (Hall, 1998) and tested the requirement of Rho family GTPases for Pk2-mediated tissue fluidity control.

We found that in the Pk2-overexpressing ectoderm, the increase in tissue fluidity was suppressed by DN-Rac1 (Fig. 8D, F, G-I) but enhanced by DN-RhoA (Fig. 8D, E, G-I). Neither DN-RhoA nor DN-Rac1 had a substantial effect on their own (Fig. 8A-C). RhoA and Rac1 have been reported to have mutually antagonistic effects on actomyosin and AJs (Burridge and Wennerberg, 2004; Chauhan et al., 2011). These observations suggest that Pk2 requires Rac1 to promote AJ remodeling.

**Figure 8.**
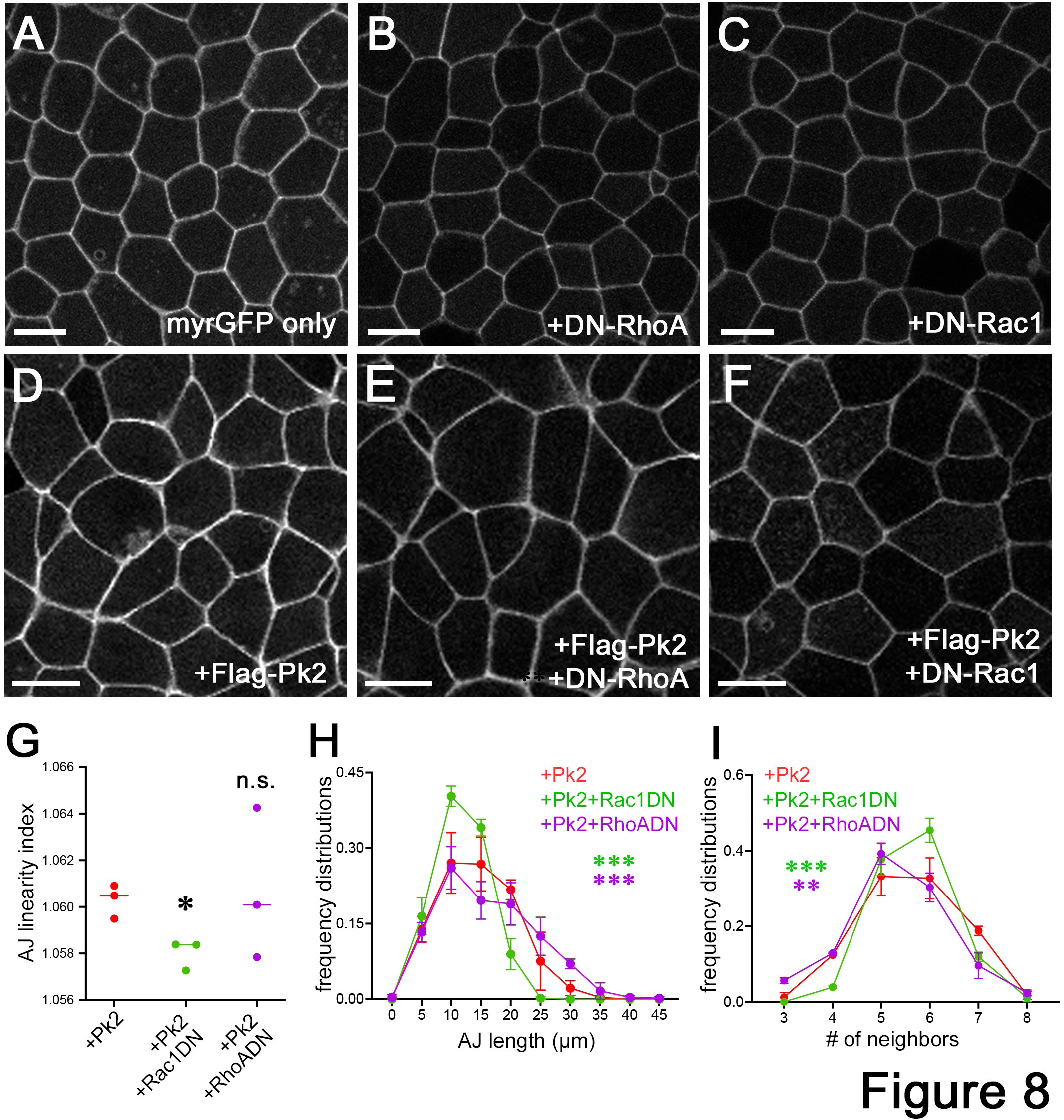
**Rac1 activation is required for the Pk2-mediated increase in tissue fluidity** (A-F) Representative images of *Xenopus* gastrula ectoderms expressing myrGFP only (A), myrGFP with DN-RhoA (B), DN-Rac1 (C), Flag-Pk2 (D), Flag-Pk2 and DN-RhoA (E), or Flag-Pk2 and DN-Rac1 (F). stage 11.5. Pigmented wild-type embryos were used for image acquisition. (G-I) Quantification of tissue fluidity according to the variation in AJ length (G), AJ linearity (H) and the number of neighboring cells (I). The numbers of cells and AJs were as follows: 291 cells and 890 AJs for Pk2; 216 cells and 655 AJs for Pk2 with DN-RhoA; and 408 cells and 1077 AJs for Pk2 with DN-Rac1. The number is the cumulative total of three embryos per group. The statistical significance of the difference between Pk2 and Pk2 with DN-RhoA or DN-Rac1 was determined. Student’s t test for G, the Kruskal‒Wallis test for H, and chi‒square test for I. **p<0.01, ***p<0.001, n.s. not significant. Scale bars: 20 μm.

### Pk2 promotes neuroepithelial cell elongation in the anteroposterior direction

While Pk2 OE promoted AJ remodeling and cell intercalation in the gastrula ectoderm, it did not have anisotropic effects (Video 5, quantification not shown). These findings suggest that factors other than Pk2 contribute to the directionality of AJ remodeling in the NE. To test this hypothesis, Pk2 RNA was overexpressed uniformly in the NE cells where Pk2 protein was distributed equally at AJs (Fig. 9A, B) similarly to its distribution in the gastrula ectoderm (Fig. S4A). The uniform Pk2 OE increased the fraction of NE cells extended along the AP axis (Fig. 9C, F, Video 9). Although Pk1 OE did not have this activity (Fig. 9D, G, Video 9), Pk1(2STR) did (Fig. 9E, H, Video 9), further supporting the critical role of the STR domain for Pk2 in directional AJ remodeling. Overall, our findings suggest that Pk2 has a permissive rather than instructive role in anisotropic AJ remodeling in the NE and requires another factor(s) for the directionality of the response.

**Figure 9.**
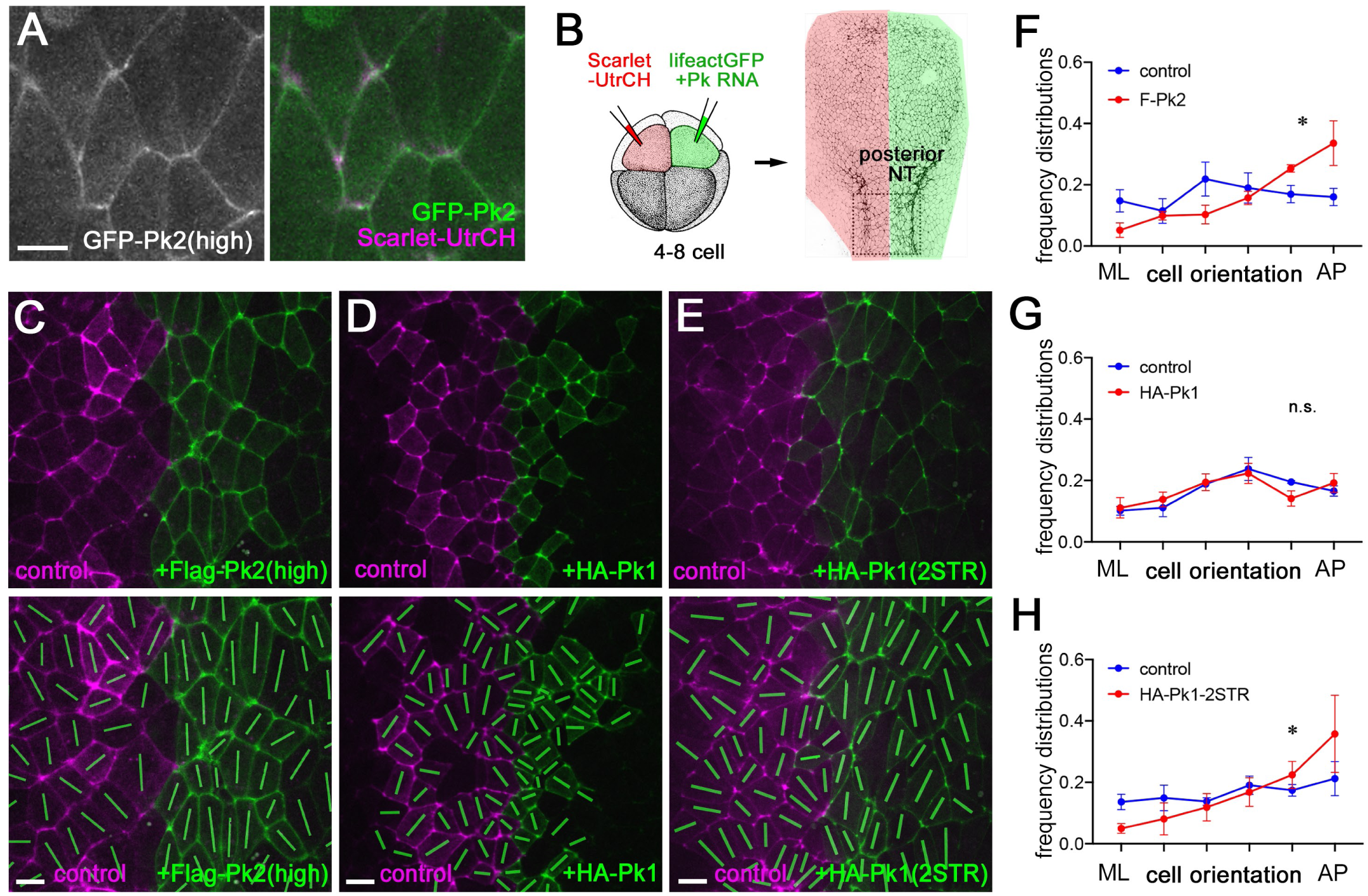
**Uniform Pk2 promotes anteroposterior elongation in neuroectoderm cells** (A) GFP-Pk2 distribution (high) in stage 14-15 NEs with Scarlet-UtrCH as an apical junction marker. (B) Scheme of the experiments for Figs. 9 and 10. LifeactGFP RNA was coinjected with HA-Pk1, Flag-Pk2 or HA-Pk1(2STR) RNA into one dorsal-animal blastomere (green) at the 4-8-cell stage. Scarlet-UtrCH RNA was injected into the other dorsal-animal blastomere (red) as a control. The posterior NE close to the border between the brain and the spinal cord was imaged. (C-E) Representative images of NEs expressing LifeactGFP with Flag-Pk2 (C), HA-Pk1 (E), or HA-Pk1(2STR) (F) from Video 9. Arrowheads indicate the presumable midline of the embryos. (F-H) The frequency distribution histogram of cell orientation in the NE cells shown in C-E. The control and Pk2 KD side of the NE in the same time-lapse imaging field were compared in three embryos per group. The number of cells per group: n=403 (control) and 562 (Flag-Pk2 (high)) (G); n=367 (control) and 350 (HA-Pk1) (H); n=484 (control) and 528 (HA-Pk1 (2STR)). Scale bars: 20 μm. The Kruskal‒Wallis test for F-H. *p<0.05.

The Vangl and Prickle family proteins preferentially localize at anterior AJs in individual NE cells (Butler and Wallingford, 2018), a hallmark of PCP. To test whether the polarized distribution of Pk2 contributes to anisotropic AJ remodeling in the NE, a low dose of Pk2 RNA was injected, at which Pk2 predominantly localized at anterior AJs and TCJs (Fig. 10A) and did not disrupt apical constriction or NTC (Fig. 10B, Video 10). The anisotropic AJ remodeling was enhanced by the low dose of Pk2 OE (Fig. 10C-E). Quantification showed that the rate of AJ elongation increased (Fig. 10F), while the rate of AJ shrinkage did not (Fig. 10G). Taken together, these results suggest that while the Pk2 protein itself does not play an instructive role *per se*, the planar localization of Pk2 at anterior AJs enhances anisotropic AJ remodeling in the NE.

**Figure 10.**
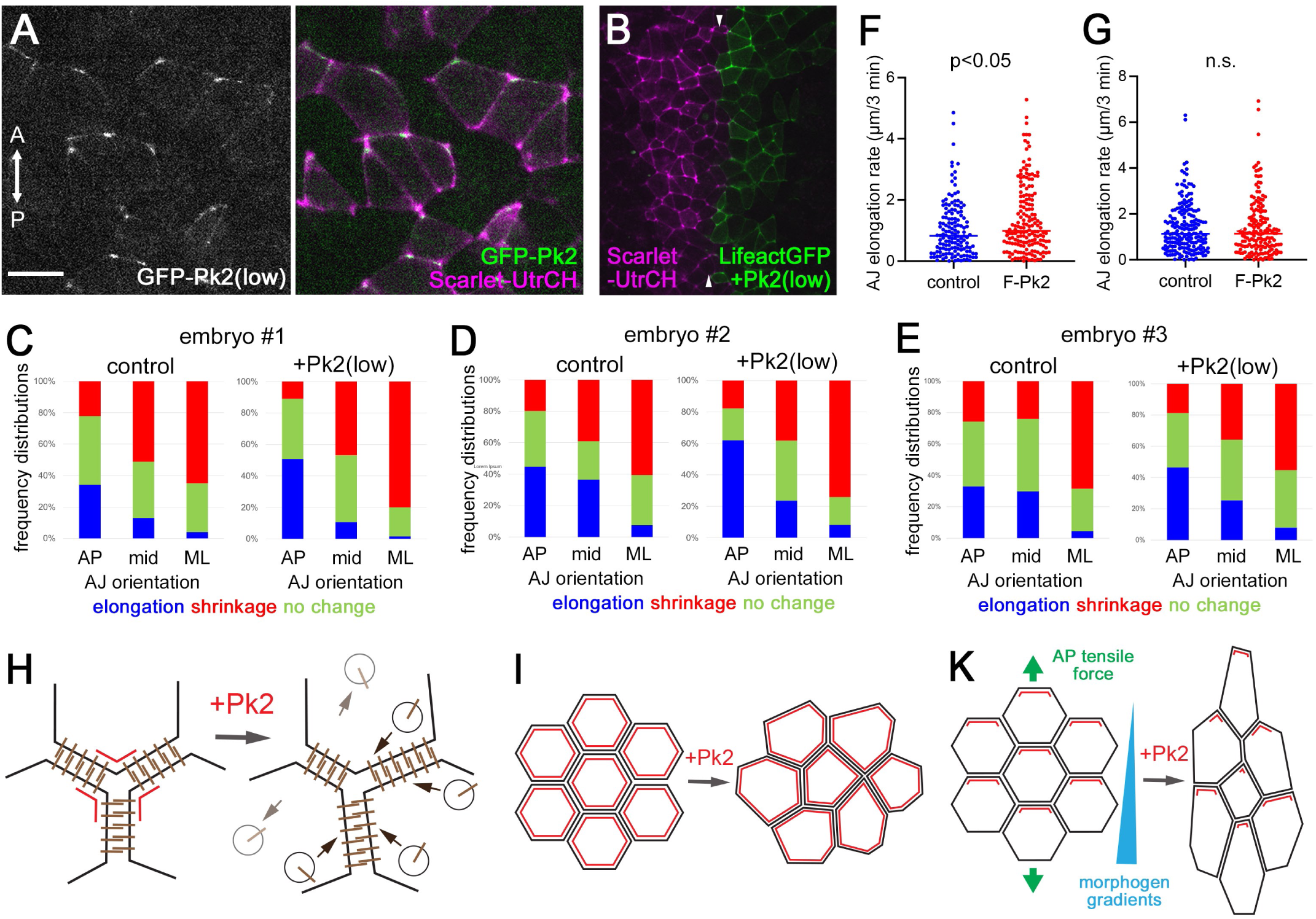
**Increased Pk2 at anterior AJs enhances anisotropic AJ remodeling in the NE** (A) Distribution of GFP-Pk2 (low dose, 50-100 pg RNA) in the NE. Scarlet-UtrCH marks apical junctions. (B) A snap-shot image of the NEs expressing control Scarlet-UtrDH only (purple, left) and Flag-Pk2 (low) with LifeactGFP (green right) from Video 10. (C-E) Quantification of changes in AJ length in the control side (left) and Flag-Pk2 side (right) of NEs in three independent embryos. Changes in AJ length are defined as follows: elongation (blue, ≥0.8 μm/3 min), shrinkage (red, ≤-0.8 μm/3 min), or no change (green, –0.8< and <0.8 μm/3 min). AJs are grouped into AP (≤30° from the AP axis), ML (≤30° from the ML axis), and mid (30-45° from either the AP or ML axes). (E, F) Rate of AJ elongation (E) and shrinkage (F). The total counts in the plots: n=291 (control) and n=280 (Pk2 (low)) for C, n=253 (control) and n=183 (Flag-Pk2) for D, and n=360 (control) and n=363 (Pk2 (low)) for E. The control and Pk2(low) NEs in the same time-lapse imaging field were compared. Scale bars: 20 μm. (I-K) Models of Pk2-mediated tissue fluidity control. Pk2 (red) increases the turnover of cadherin-based AJs by influencing the components of cell-cell adhesion complexes or the underlying actomyosin networks, promoting AJ remodeling. This is important to increase the susceptibility of AJs to stimuli-induced remodeling (I). In the absence of directional cues, such as in the gastrula ectoderm, Pk2 increases nondirectional AJ remodeling (J). In the NE, AP-oriented cues in the environment, such as AP tensile force and morphogen gradients, and planar polarization of Pk2 cooperatively promote anisotropic AJ remodeling (K).

## Discussion

This study reveals a previously uncharacterized role for Pk2 in tissue fluidity control in the *Xenopus* ectoderm during neurulation. We show that Pk2 OE increased the rate of AJ elongation and shrinking in non-neural ectoderm. In the NE, the rate of junction elongation was also increased by Pk2 OE. Conversely, the rate of both junction elongation and shrinking was reduced in the Pk2 KD. Thus, Pk2 increases the efficiency of AJ elongation and shrinking to stimulate cell rearrangement and epithelial deformation (Fig. 10H, I). Although non-neural ectoderm isotropically reacted to Pk2, the effect of Pk2 on AJs in the NE was anisotropic. In the wild-type NE, AP junctions tend to elongate, whereas ML junctions shrink. Pk2 OE increased the frequencies of elongating AP and shrinking ML junctions. Pk2-depleted NE had a complementary phenotype: the frequencies of both elongating AP junctions and shrinking ML junctions decreased, resulting in defective NTC. We propose that the observed directionality of the response is due to chemical or mechanical cues in the NE that cooperate with Pk2 to instruct morphogenesis, acting preferentially along the AP axis (Fig. 10K).

Given the anterior enrichment of Pk2 and Vangl2 at anterior bicellular AJs (Butler and Wallingford, 2018; Ossipova et al., 2015c) and TCJs (Fig. 10A), Pk2 may locally increase the efficiency of anterior junction remodeling by modulating cell adhesion or actomyosin networks (Fig. 10H). Although the Vangl2 binding domain is required for Pk2-induced AJ remodeling, the specific role of Vangl2 in tissue fluidity remains unclear, because neither Vangl2 overexpression nor knockdown mimic the observed Pk2 effects on AJs (Matsuda et al., 2023). We propose that Pk2 planar polarization amplifies the effect of environmental cues on AJ remodeling along the AP axis (Fig. 10K). These cues may include tensile stresses originating from the AP elongation of embryo body axis (Christodoulou and Skourides, 2022; Clausi and Brodland, 1993; Davidson and Keller, 1999; Handler et al., 2023; Hirano et al., 2022; Moon and Xiong, 2022; Smith and Schoenwolf, 1997; Sokol, 2016; Xiong et al., 2020; Zhou et al., 2015). Alternatively, diffusible morphogen gradients may be responsible the directionality of Pk2 effects. Supporting the latter possibility, several Wnt and FGF ligands are expressed predominantly posteriorly, in a graded manner during neurulation, while retinoic acid (RA) establishes a reverse gradient (Aulehla and Pourquie, 2010; Diez Del Corral and Morales, 2017; Edri et al., 2023; Niederreither et al., 1999; Nordstrom et al., 2002; Storey et al., 1998; Yamaguchi, 2001; Yoon et al., 2023).

Compared with other vertebrate Pk proteins, the ability of Pk2 to remodel AJs appears to be unique. A specific region in the C-terminal region of Pk2 (STR) appears to be responsible for the functional differences between Pk1 and Pk2, because it conferred the AJ remodeling activity of Pk2 to Pk1. Whereas *Drosophila* Prickle lacks the STR region, its presence in Pk1 and Pk2 parallels the ability of the vertebrate PCP pathway to influence morphogenesis. Notably, Pk1 was shown to modulate F-*actin* both *in vivo* and *in vitro* (Carreira-Barbosa et al., 2003; Huang and Winklbauer, 2022; Tao et al., 2009; Veeman et al., 2003). In our model, a regulator of Rho GTPases binds to Pk2 via its STR domain and locally activates Rac1, increasing the efficiency of AJ remodeling. The identification of these putative GAPs or GEFs will be important for further elucidating the underlying mechanism of Pk2-mediated AJ remodeling.

## Methods

### Plasmids, morpholino and RNA preparation

pCS107-GFP-xPk2 was kindly provided by J. Wallingford (University of Texas)(Butler and Wallingford, 2015; Butler and Wallingford, 2018). xPk1S cDNA was purchased from Open Biosystems (Horizon). pCS2-3xFlag-xPk3 and pCS2-GFP-xPk3 was generated from pCS2-Flag-Pk3 (Ossipova et al., 2015a) by PCR using primers including the correct C-terminal sequence. Subcloned Pk2 deletion mutants or Pk1S were inserted into either pCS107-Flag or pCS2-3xHA using restriction enzymes and T4 ligase. The domains in the C-terminal half of Pk2 are defined as follows: C1 (aa. 367-561), STR(aa. 562-665), C4(aa. 666-733) and VBD (aa. 734-827). The NEBuilder HiFi DNA Assembly Kit (NEB) was used for the construction of chimeric Pk1 and Pk2 mutants. pCS2-GFP, myrGFP and myrRFP were described previously (Matsuda et al., 2023). pCS2-lifeactGFP was constructed from pCS2-GFP and annealed to two oligonucleotides encoding Lifeact probe peptides. Scarlet-UtrCH was subcloned and inserted into the pCS2 vector from 3xmScarlet I_UtrCH, a gift from Dorus Gadella (Addgene plasmid # 112962). pT7T-C-cadherin-mYFP was modified from pT7T-CcadTSMod, a gift from Marc Tramier (Addgene plasmid # 99874)(Herbomel et al., 2017). GFP-αE-catenin was subcloned and inserted into pCS2 from pCAGGS-Flag-αE-catenin, kindly provided by Yoshimi Takai (Kobe University, Japan)(Sakakibara et al., 2020). pCS2-mNeon-Sf9 and pCS2-GFP-tricellulin were generous gifts from Ed Munro (University of Chicago, USA)(Hashimoto et al., 2015) and Ann Miller (University of Michigan, USA)(Higashi et al., 2016). DN-RhoA and DN-Rac1 in pRK5 were gifts from Alan Hall (Sloan-Kettering Institute, USA)(Hall, 1998).

Capped RNA was synthesized using an mMessage mMachine SP6 Transcription Kit (Invitrogen) and purified with RNeasy Mini Kit (Qiagen). *Pk2* splicing blocking morpholino (MO) (5’– GAACCCAAACAAACACTTACCTGTT –3’) and *Pk1* splicing blocking MO (5’– CTTCTGATCCATTTCCAAAGGCATG –3’) (GeneTools) were characterized in previous studies (Butler and Wallingford, 2018; Daulat et al., 2012; Huang and Winklbauer, 2022; Takeuchi et al., 2003).

### Xenopus embryos and microinjection

Wild-type and albino *Xenopus laevis* were purchased from Nasco and Xenopus1 and were maintained and handled following the Guide for the Care and Use of Laboratory Animals of the National Institutes of Health. The protocol for animal use was approved by the Institutional Animal Care and Use Committee (IACUC) of the Icahn School of Medicine at Mount Sinai. The sex of the animals was not considered in the study design and analysis because the study subjects were sexually indifferent embryos. In vitro fertilization and embryo culture were performed as previously described (Ossipova et al., 2014). Embryo staging was determined according to (Nieuwkoop and Faber, 1994). For microinjections, embryos were transferred to 3% Ficoll 400 (Pharmacia) in 0.5× Marc’s modified Ringer’s (MMR) solution (50 mM NaCl, 1 mM KCl, 1 mM CaCl2, 0.5 mM MgCl2 and 2.5 mM HEPES (pH 7.4))(Peng, 1991).

RNA or MO in 5-10 nl of RNase-free water (Invitrogen) was microinjected into one to two animal blastomeres of 4-8 cell stage embryos. For apical domain imaging of gastrula embryos, two ventral-animal blastomeres were injected with 100 pg of RNA encoding Flag– or HA-tagged Pk proteins and coinjected with 50 pg of myrGFP RNA. For apical domain imaging of the neurula, one dorsal-animal blastomere was injected with Pk2 morpholino (MO) or RNA encoding 100 pg of Flag– or HA-tagged Pks, together with 50 pg of myrGFP or LifeactGFP RNA. Subsequently, the other dorsal-animal blastomere in the same embryos was injected with RNA encoding 50 pg of myrRFP or Scarlet-UtrCH. For imaging to determine Pk2 distribution, RNA encoding GFP-Pk2 was coinjected with RNA encoding myrRFP or Scarlet-UtrCH. RNA– or MO-injected embryos were cultured in 0.1x MMR until the early gastrula or neurula stage. Each injection included at least 20 embryos per condition. The experiments were repeated at least three times.

### Phalloidin staining and imaging of fixed Xenopus embryos

For phalloidin staining of the neurula, embryos were fixed in MEMFA (100 mM MOPS (pH 7.4), 2 mM EGTA, 1 mM MgSO_4_, 3.7% formaldehyde)(Harland, 1991) for 1 hr at room temperature. After permeabilization in 0.1% Triton X-100 in PBS for 10 min, the embryos were incubated with Alexa Fluor 555-conjugated phalloidin (1:400 dilution, Thermo Fisher Scientific) in PBS containing 1% BSA overnight at 4°C. The dissected neural plate was mounted on a glass slide with two coverglass spacers (0.13-0.17 mm) to minimize damage to the morphology.

Images of the fixed embryos were captured either by the BC43 spinning disk confocal microscope (Fusion Ver 2, Andor, Oxford Instruments) or the Zeiss LSM 980 with Airyscan 2 confocal microscope (Zen (blue edition) Ver 3.7.97.04000, Zeiss). Either Nikon CFI Plan Apochromat Lambda D 20X (NA=0.8 and WD=0.8 mm), Nikon CFI Plan Apo Lambda S 40XC Sil (NA=1.25 and WD=0.3 mm), Zeiss Plan-Apochromat 20x/0.8 (NA=0.8 and WD=0.55 mm) or Zeiss C-Apochromat 40x/1.2 W (NA=1.2 and WD=0.28 mm) was used for gastrula imaging. Nikon CFI Plan Apochromat Lambda D 20X (NA=0.8 and WD=0.8 mm) was used for the neurula stage. The experiments were repeated at least three times. Each imaging included at least 3 embryos per experimental group. Z-stack images were projected into a single image using the maximum projection function in ImageJ2 (ver. 2.9.0/1.54g), which was used for further analysis and quantification.

### Time-lapse imaging of Xenopus embryos

Embryos were injected with RNA encoding Pks and fluorescent protein-tagged marker proteins (myrGFP, myrRFP, LifeactGFP, 3xScarlet-UtrCH, GFP-tricellulin, mNeon-Sf9, YFP-C-cadherin, and GFP-αE-catenin). Gastrula or neurula were mounted in 1% low melting temperature agarose (SeaPlaque agarose, #50101, Lonza) in 0.1× MMR on a glass slide attached to a silicone isolator (1.2 mm, Grace Biolabs) or on a glass-bottom dish (#1, Cellvis). Time-lapse imaging was carried out using a Andor BC43 spinning disk confocal microscope or a Zeiss LSM980 confocal microscope, or a Zeiss AxioZoomV16 fluorescence stereomicroscope equipped with an AxioCam 506 mono CCD camera (Zen (blue edition), Zeiss) at room temperature. Images were taken every 3-6 min for a period of 0.5-5 hrs. The multiposition function in imaging softwares was used for simultaneous time-lapse imaging of embryos in each experimental group. Z-stack images were projected into a single image using the maximum projection function in ImageJ.

### Data analysis, segmentation, cell tracking, and apical domain assessment

Image processing and quantification were performed as previously described (Matsuda et al., 2023). Briefly, grayscale images of the cell outline marker were segmented using the Python package Cellpose v2.0.5 (Pachitariu and Stringer, 2022). Segmented cells were tracked across timepoints using the Python package Bayesian Tracker (btrack v0.4.5)(Ulicna et al., 2021) with manual correction. For NP segmentation, the NP area in stage 14-15 embryos was defined either by brighter signals of LifeactGFP, Scarlet-UtrCH or phalloidin staining than in the surrounding epidermis. Pigmented wild-type embryos were used for image segmentation.

### Analysis of junctional morphology and fluorescence intensities

The morphology and fluorescence intensity of individual cell junctions were quantified as described previously (Matsuda et al., 2023). Briefly, after segmentation and tracking, each junction was first identified as a vertex or edge of the cell outline network, and mean protein intensities were measured using dilated masks of the pixel or a set of pixels that defined the junction.

### Statistics & Reproducibility

Histograms and dot plots of the experimental data were generated using GraphPad Prism 10 (ver. 10.1.0) and Microsoft Excel. Individual experiments were repeated at least three times. The coefficient of variation (CV) and s.d. were calculated using GraphPad Prism 10 and used to measure the variability in individual sample groups. Student’s t test or the Kolmogorov‒Smirnov test was used to determine the statistical significance of differences between two groups with normal or nonnormal distributions. Chi-square test or the Kruskal‒Wallis test was used for the categorized or ranked data.

## Acknowledgments

We thank D. Gadella, A. Hall, A. Miller, E. Munro, Y. Takai, M. Tramier and J. Wallingford, for plasmids. We also thank Jakub Harnos and Chih-Wen Chu for the construction of plasmids encoding Pk2 deletion mutants. We are grateful to Tanya Whitfield, Karen Kasza and Sassan Ostwar for comments on the manuscript and members of the Sokol laboratory for valuable discussions. We acknowledge the help from the ISMMS Microscopy Core facility. This research was supported by the NIH grant R35GM122492 to S.Y.S.

## Author contribution statement

M.M. and S.Y.S. conceptualized and developed the project. M.M. designed and performed the experiments and the formal analyses. M.M. and S.Y.S. contributed to the data interpretation. M.M. prepared the figures and wrote the original manuscript. M.M. and S.Y.S. edited the manuscript. Both authors approved the final manuscript.

## Supplemental information

**Supplemental Figure 1 Pk2 knockdown inhibits anteroposterior cell orientation in the neuroectoderm**

(A) Scheme of the experiment. Pk2 MO and GFP RNA were coinjected into one dorsal-animal blastomere (green) at the 4-8-cell stage. Fixed embryos (stage 15) were stained with phalloidin. The posterior neural plate near the brain‒spinal cord border was imaged. (B-B’”) Representative images of Pk2-KD NEs. Phalloidin staining only (B), overlayed color coding based on apical domain size (B’), green bars indicating cell orientation (B”), and GFP-labeled Pk2 MO cells (B’”). (C) Frequency distribution histogram of cell orientation in the control (blue) or Pk2 KD (red) NEs. The control and Pk2 KD side of the NE in the same imaging field were compared in three embryos per group. The means +/− SEMs are shown. n=920 (control) and n=1093 (Pk2 KD). Four embryos per group. (D) 2D plots of AJ orientation (x-axis) and length (y-axis) in the control NEs (left) and Pk2-KD NEs (right). Plots include the data obtained from tracking 24 AJs (control) and 16 AJs (Pk2 KD) from a time-lapse movie with total durations of 141 min and 3 min intervals. The total counts in the plots were 377 (control) and 302 (Pk2 MO). (E) Quantification of changes in the length of AJs in control NEs (left) and Pk2 MO NEs (right). Changes in AJ length are defined as follows: elongation (blue, >0.8 μm/3 min), shrinkage (red, >0.8 μm/3 min), or no change (green, <0.8 μm/3 min). AJs are grouped into AP (≤30° from the AP axis), ML (≤30° from the ML axis), and mid (30-45° from either the AP or ML axes). The control and Pk2 KD side of the NE in the same time-lapse imaging field were compared. Chi‒square test for E.

**Supplemental Figure 2. Pk2 promotes the elongation of newly formed AJs after cytokinesis**

(A, B) Representative images of newly formed AJs after cytokinesis. *Xenopus* gastrula ectoderm expressing GFP-ZO1 only (A) or with Flag-Pk2 (B). Time-lapse imaging was initiated at stage 11, and t=00:00 was set at the time when the contractile ring was closed. (C) AJ length changes in cells expressing GFP-ZO1 only (blue) or with HA-RFP-Pk2 (red). t=00:00 was set at the time when the contractile ring was closed. Three embryos per group. The number of newly formed AJs: n=38 (control); n=20 (HA-RFP-Pk2). The statistical significance of differences was calculated based on the maximum length of the AJs after cytokinesis. p<0.0001. Student’s t test. Scale bars: 10 μm.

**Supplemental Figure 3. Pk2 reduces AJ and actomyosin network components at TCJs**

(A) Representative images of Sf9-mNeon in control or Flag-Pk2-expressing gastrula ectoderm, stage 11.5. (B) Representative images of YFP-αE-catenin in control (left) or Pk2 (right) gastrula ectoderms. stage 11.5. (C, D) Quantification of the fluorescence intensity of mNeon-Sf9 (C) or YFP-αE-catenin (D) at TCJs. The fluorescence intensity at TCJs was divided by the average fluorescence intensity at BJs. Total number of TCJs: n=59 (control) and n=54 (Pk2) for mNeon-Sf9; n=64 (control) and n=53 (Pk2) for YFP-αE-catenin. The Kolmogorov-Smirnov test. *p<0.05, **p<0.01. Scale bars: 10 μm.

**Supplemental Figure 4. Subcellular distribution of GFP-Pk2 mutant proteins in the gastrula ectoderm**

Representative images of gastrula ectoderm cells expressing GFP-Pk2 (A), GFP-Pk2ΔSTR (B), GFP-Pk2ΔC4 (C), GFP-Pk2ΔVBD (D) or GFP-Pk2ΔC30 (E). Stage 11. myrRFP RNA was coinjected to mark the plasma membrane. Scale bars: 20 μm.

**Supplemental Figure 5. Neither Pk1 nor Pk3 increases tissue fluidity in the gastrula ectoderm**

(A) Schematic diagrams of Pk1, Pk2 and Pk3. Pk3 does not contain the conserved STR (blue) or C4 (light green) regions. (B) Alignment of the STR region of human, mouse, *Xenopus* and *zebrafish* Pk1 and Pk2. The conserved amino acids among Pk1 and Pk2 (red), only among Pk1 (blue) or only among Pk2 (green).(C-I) Representative images of *Xenopus* gastrula ectoderms expressing myrGFP only (C, F), myrGFP with HA-Pk1 (D), Flag-Pk2 (E), Flag-Pk3 (G), HA-Vangl2 (H), or Flag-Pk3 and HA-Vangl2 (I). stage 11.5. Pigmented wild-type embryos were used. (J-L) Quantification of tissue fluidity according to the variation in AJ length (J), AJ linearity (K) and the number of neighboring cells (L). A total of 281 cells and 827 AJs were scored in the control group (myrGFP only); 191 cells and 578 AJs were scored in the Pk3 group; 329 cells and 959 AJs were scored in the Vangl2 group; and 368 cells and 1068 AJs were scored in the control group (Pk3 and Vangl2) in J-K. Three embryos per group. Student’s t test for J, the Kruskal‒Wallis test for K, and chi‒square test for L. n.s., not significant. Scale bars: 10 μm.

## Supplemental Videos

**Video 1.** A representative time-lapse Video showing the dynamics of NE cells during NTC in Pk2 KD embryos. Pk2 MO was coinjected with RNA encoding LifeactGFP in one dorsal blastomere at the 4-8-cell stage and RNA encoding Scarlet-UtrCH was injected into the other dorsal blastomere. The posterior NE near the border between the head and the spinal cord was imaged from the dorsal side (dorsal view as shown in Fig. 1A). Time-lapse imaging was initiated at stage 14. 5 min interval. The time-lapse duration was 3 hrs 40 min at room temperature.

**Video 2.** A representative time-lapse video of the control (left) and Pk2-depleted NEs (right). Scarlet-UtrCH was injected into one dorsal blastomere. Pk2 MO was coinjected with LifeactGFP in the other dorsal blastomere. Dorsal view. Stage 14. 3 min interval. Time-lapse duration: 1 hr 12 min at room temperature.

**Video 3.** A representative time-lapse video showing changes in cell orientation (green lines) in control NEs and Pk2 KD NEs. Pk2 MO was coinjected with RNA encoding myrGFP in one dorsal blastomere at the 4-8-cell stage. RNA encoding myrRFP was injected into the other dorsal blastomere. Dorsal view. Stage 15. 3 min interval. The time-lapse duration was 1 h 12 min at room temperature.

**Video 4.** A representative time-lapse video showing the dynamics of NE cells during NTC in Pk1 KD embryos. Pk1 MO was coinjected with RNA encoding LifeactGFP in one dorsal blastomere at the 4-8-cell stage, and RNA encoding Scarlet-UtrCH was injected into the other dorsal blastomere. Dorsal view. Stage 15. 5 min interval. The time-lapse duration was 2 hours 55 min at room temperature.

**Video 5.** A representative time-lapse video of the Pk1 morphant NE. Pk1MO was coinjected with LifeactGFP in one dorsal blastomere. Dorsal view. Stage 14. 5 min interval. The time-lapse duration was 1 hr 45 min at room temperature.

**Video 6.** A representative time-lapse video showing hexagonal packing during gastrulation in wild-type (left) and Flag-Pk2 OE (right) embryos. RNA encoding myrGFP was injected with or without Flag-Pk2 RNA into two ventral blastomeres of 4-cell-stage embryos. The animal side ectoderm was imaged from the top of the embryos, albino embryos were used. Time-lapse imaging was initiated at stage 10. 5 min interval. The time-lapse duration was 4 hrs 55 min at room temperature.

**Video 7.** Representative time-lapse video showing hexagonal packing and the lateral domain dynamics in the control wild-type (left) and Flag-Pk2 OE (right) ectoderms. Injection was performed as described in Fig. 3A. The animal side ectoderm was imaged. The maximum intensity projection (Image J) was applied to z-stack images. Stage 11.5, albino embryos. 5 min interval. Time-lapse duration: 1 hour 45 min at room temperature.

**Video 8.** A representative time-lapse video showing YFP-C-cadherin dynamics in the control wild-type (left) and Flag-Pk2 OE (right) ectoderms. RNA encoding C-cadherin-YFP was injected into two ventral blastomeres of 4-cell-stage embryos with or without RNA encoding Flag-Pk2. The animal side ectoderm was imaged. Time-lapse imaging was initiated at stage 11. 30-second intervals. Time-lapse duration: 14 min 30 sec at room temperature.

**Video 9.** A representative time-lapse video showing the dynamics of NE cells expressing high levels of Flag-Pk2 (left panel), HA-Pk1 (middle panel) or HA-Pk1(2STR) (right panel). RNA encoding Scarlet-UtrCH was injected into one dorsal blastomere of 4-cell-stage embryos (purple on the left side of the NE). RNAs encoding LifeactGFP and various Pk constructs were coinjected into the other dorsal blastomere. Dorsal view. Time-lapse imaging was initiated at stage 14. 6 min interval. The time-lapse duration was 2 hrs 36 min at room temperature.

**Video 10.** A representative time-lapse video showing the dynamics of NE cells and NTC in embryos expressing low levels of Pk2. 100 pg of RNA encoding Flag-Pk2 was coinjected with LifeactGFP in one dorsal blastomere. RNA encoding Scarlet-UtrCH was injected into the other dorsal blastomere. Dorsal view. Stage 15. 6 min interval. The time-lapse duration was 1 hr and 54 min at room temperature.

