## Supplemental Figures for "Prickle2 regulates apical junction remodeling and tissue fluidity during vertebrate neurulation"

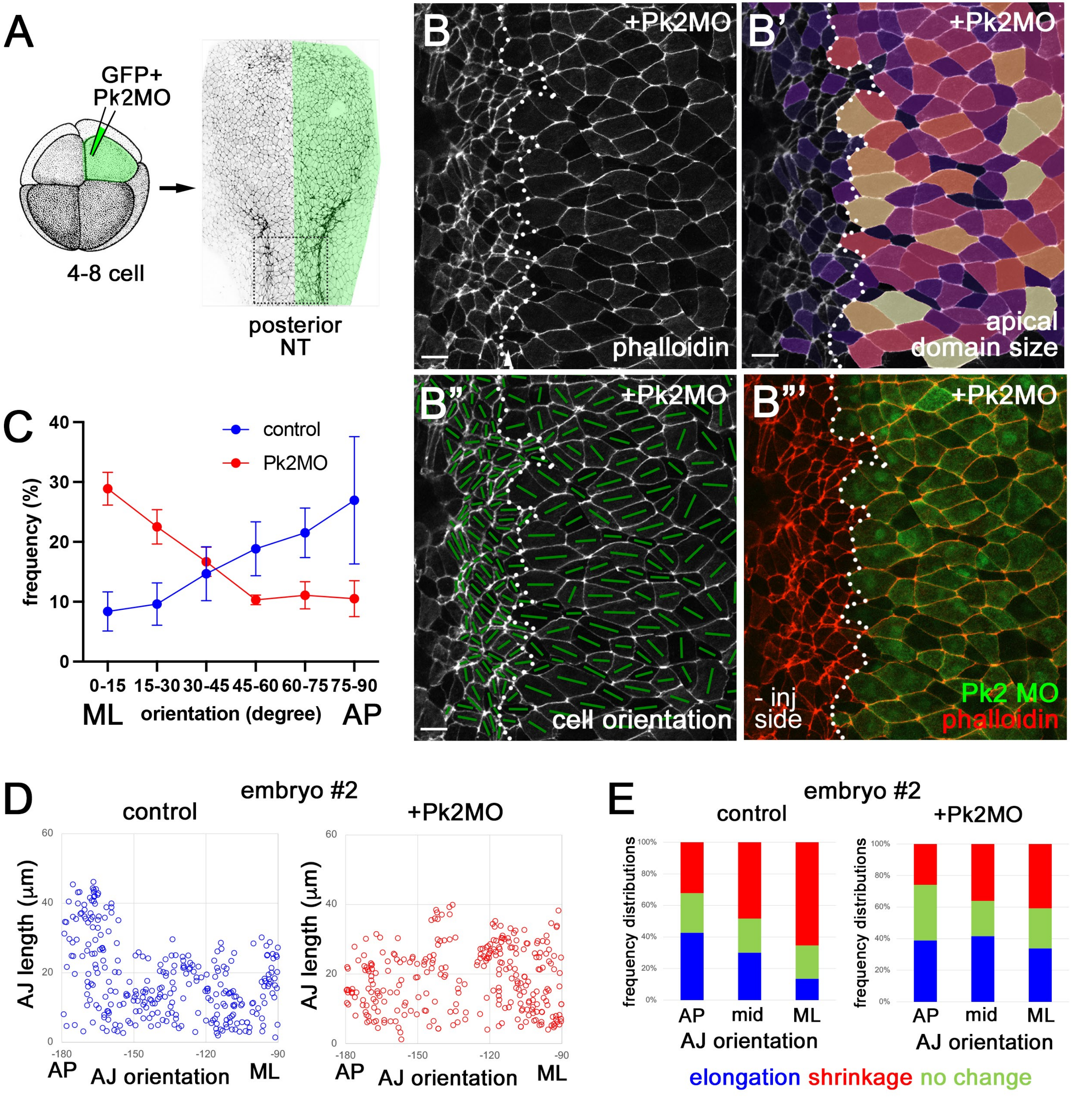
A
GFP+
Pk2MO
4-8 cell
posterior
NT
B
+Pk2MO
phalloidin
B'
+Pk2MO
apical
domain size
B"
+Pk2MO
cell orientation
B"'
+Pk2MO
- inj
side
Pk2 MO
phalloidin
C
control
Pk2MO
frequency (%)
0-15
15-30
30-45
45-60
60-75
75-90
ML
orientation (degree)
AP
D
embryo #2
control
+Pk2MO
AJ length (μm)
AP
AJ orientation
ML
E
embryo #2
control
+Pk2MO
frequency distributions
AP
mid
ML
AJ orientation
elongation shrinkage no change

### Figure S1

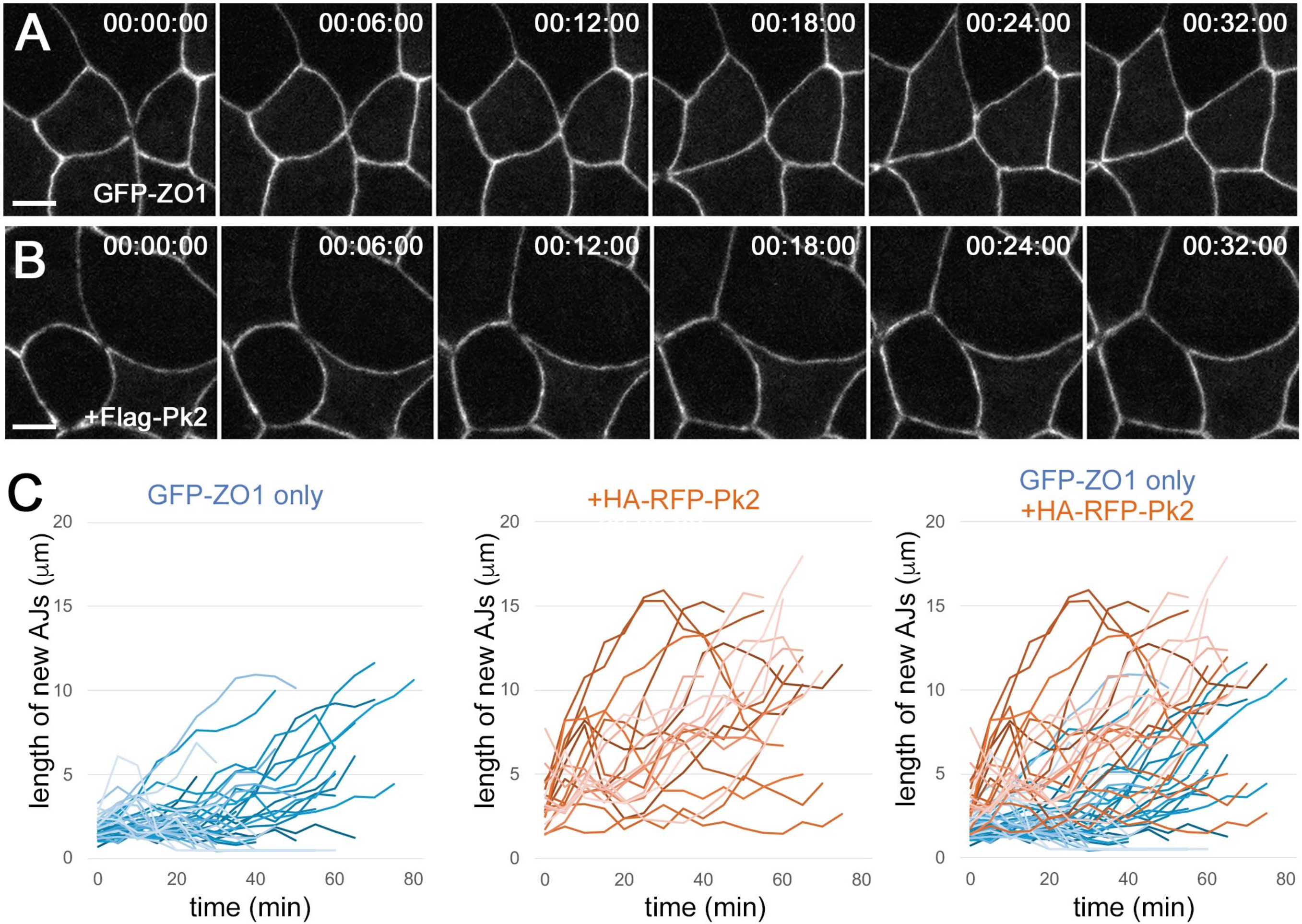
A
00:00:00
00:06:00
00:12:00
00:18:00
00:24:00
00:32:00
GFP-ZO1
B
00:00:00
00:06:00
00:12:00
00:18:00
00:24:00
00:32:00
+Flag-Pk2
C
GFP-ZO1 only
+HA-RFP-Pk2
GFP-ZO1 only
+HA-RFP-Pk2
length of new AJs (μm)
time (min)

Figure S2

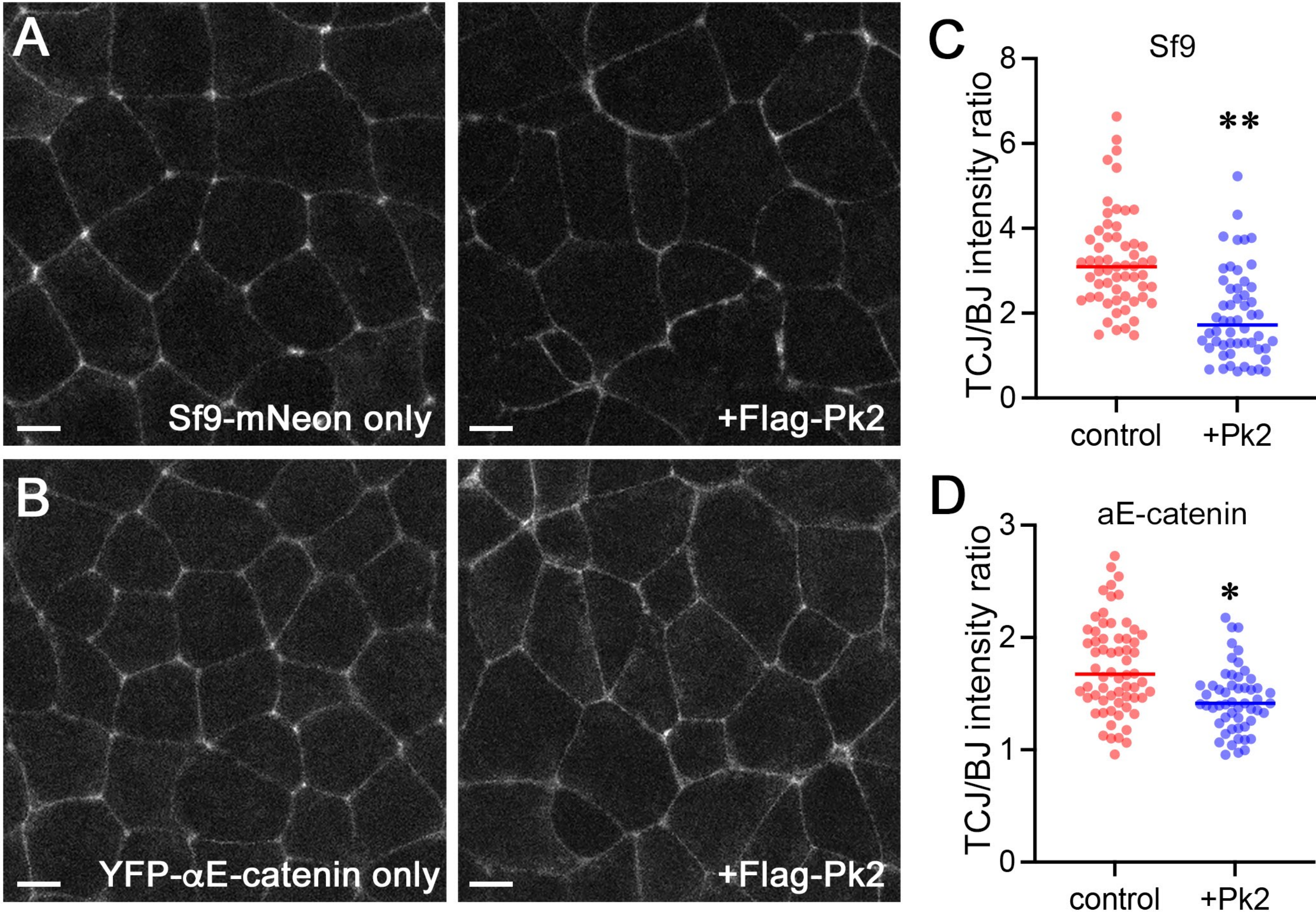
A
Sf9-mNeon only
+Flag-Pk2
B
YFP-αE-catenin only
+Flag-Pk2
C
Sf9
TCJ/BJ intensity ratio
8
6
4
2
0
**
control
+Pk2
D
aE-catenin
TCJ/BJ intensity ratio
3
2
1
0
*
control
+Pk2

### Figure S3

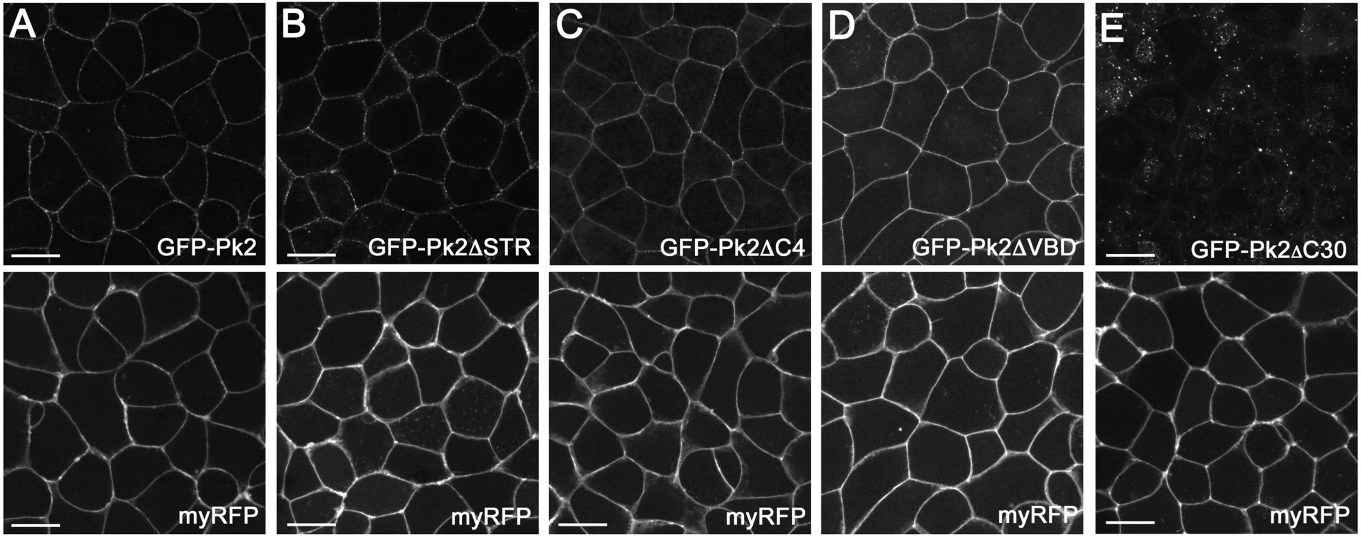
A
GFP-Pk2
B
GFP-Pk2ΔSTR
C
GFP-Pk2ΔC4
D
GFP-Pk2ΔVBD
E
GFP-Pk2ΔC30
myRFP
myRFP
myRFP
myRFP
myRFP

### Figure S4

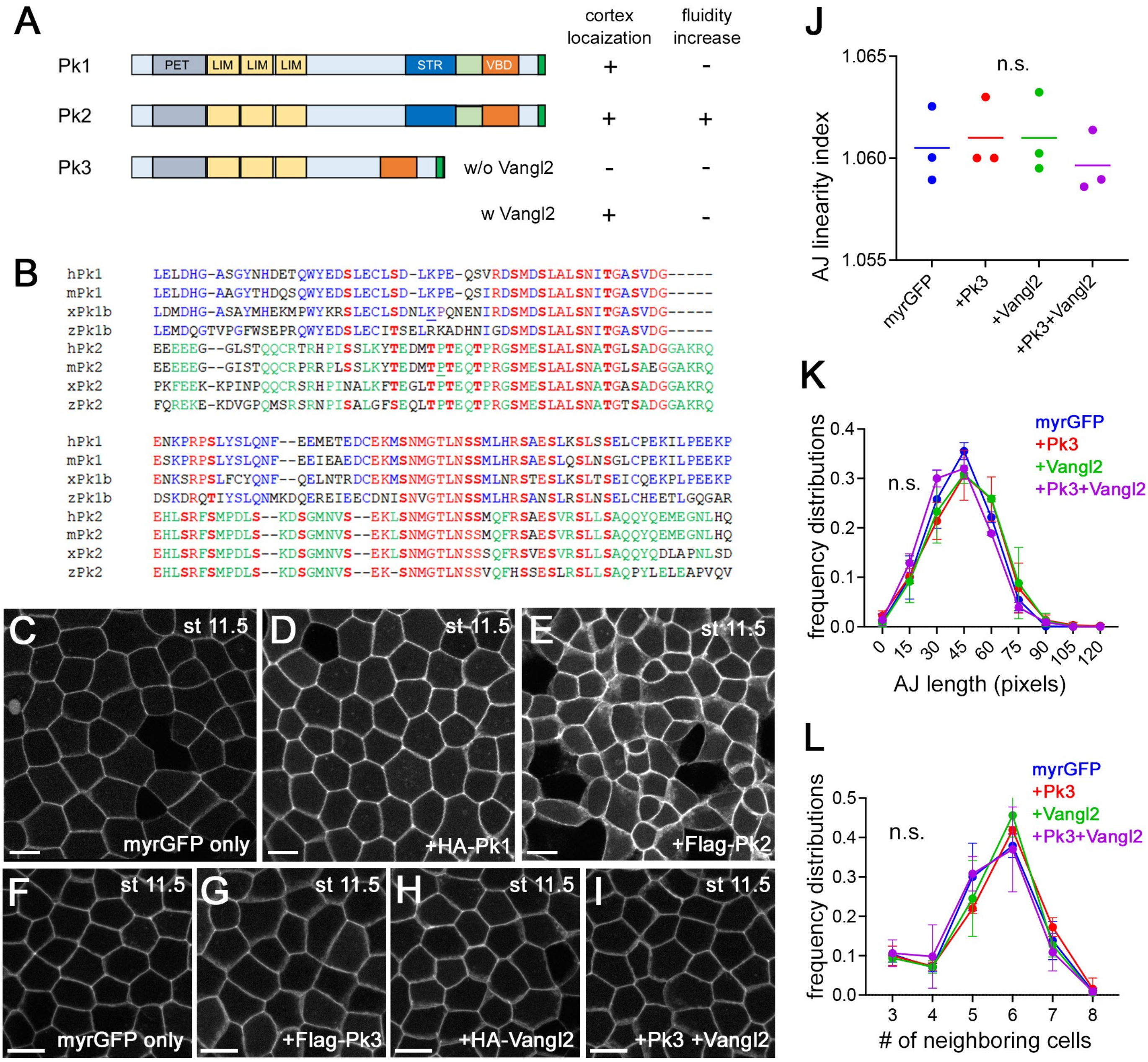

A
cortex locaization
fluidity increase
Pk1
PET
LIM
LIM
LIM
STR
VBD
+
-
Pk2
+
+
Pk3
w/o Vangl2
-
-
w Vangl2
+
-
B
hPk1 LELDHG-ASGYNHDETQWYEDSLECLSD-LKPE-QSVRDSMDSLALSNITGASVDG-----
mPk1 LELDHG-AAGYTHDQSQWYEDSLECLSD-LKPE-QSIRDSMDSLALSNITGASVDG-----
xPk1b LDMDHG-ASAYMHEKMPWYKRSLECLSDNLKPQNENIRDSMDSLALSNITGASVDG-----
zPk1b LEMDQGTVPGFWSEPRQWYEDSLECITSELRKADHNIGDSMDSLALSNITGASVDG-----
hPk2 EEEEEG--GLSTQQCRTRHPISSLKYTEDMTPTEQTPRGSMESLALSNATGLSADGGAKRQ
mPk2 EEEEEG--GISTQQCRPRRPLSSLKYTEDMTPTEQTPRGSMESLALSNATGLSAEGGAKRQ
xPk2 PKFEEK-KPINPQQCRSRHPINALKFTEGLTPTEQTPRGSMESLALSNATGASADGGAKRQ
zPk2 FQREKE-KDVGPQMSRSRNPISALGFSEQLTPTEQTPRGSMESLALSNATGTSADGGAKRQ
hPk1 ENKPRPSLYSLQNF--EEMETEDCEKMSNMGTLNSSMLHRSAESLKSLSSELCPEKILPEEKP
mPk1 ESKPRPSLYSLQNF--EEIEAEDCEKMSNMGTLNSSMLHRSAESLQSLNSGLCPEKILPEEKP
xPk1b ENKSRPSLFCYQNF--QELNTRDCEKMSNMGTLNSSMLNRSTESLKSLTSEICQEKPLPEEKP
zPk1b DSKDRQTIYSLQNMKDQEREIEECDNISNVGTLNSSMLHRSANSLRSLNSELCHEETLGQGAR
hPk2 EHLSRFSMPDLS--KDSGMNVS--EKLSNMGTLNSSMQFRSAESVRSLLSAQQYQEMEGNLHQ
mPk2 EHLSRFSMPDLS--KDSGMNVS--EKLSNMGTLNSSMQFRSAESVRSLLSAQQYQEMEGNLHQ
xPk2 EHLSRFSMPDLS--KDSGMNVS--EKLSNMGTLNSSSQFRSVESVRSLLSAQQYQDLAPNLSD
zPk2 EHLSRFSMPDLS--KDSGMNVS--EK-SNMGTLNSSVQFHSSESLRSLLSAQPYLELEAPVQV
C
st 11.5
myrGFP only
D
st 11.5
+HA-Pk1
E
st 11.5
+Flag-Pk2
F
st 11.5
myrGFP only
G
st 11.5
+Flag-Pk3
H
st 11.5
+HA-Vangl2
I
st 11.5
+Pk3 +Vangl2
J
AJ linearity index
1.065
1.060
1.055
n.s.
myrGFP
+Pk3
+Vangl2
+Pk3+Vangl2
K
frequency distributions
0.4
0.3
0.2
0.1
0.0
n.s.
myrGFP
+Pk3
+Vangl2
+Pk3+Vangl2
0
15
30
45
60
75
90
105
120
AJ length (pixels)
L
frequency distributions
0.5
0.4
0.3
0.2
0.1
0.0
n.s.
myrGFP
+Pk3
+Vangl2
+Pk3+Vangl2
3
4
5
6
7
8
### of neighboring cells
